# Genome-wide characterization of host factors involved in single-stranded RNA and DNA phage infection pathways

**DOI:** 10.64898/2026.09.17.752499

**Authors:** Isabella Murray, Roniya Thapa Magar, Denish Piya, Hemaa Selvakumar, Lucas Moriniere, Sarshad Koderi Valappil, Madeline Svab, Jirapat Thongchol, Alexey Kazakov, Nathalie H. Elisabeth, Mohamad Alayouni, Prithvi Pal Singh, Artur Muszyński, Parastoo Azadi, Siân V. Owen, Junjie Zhang, Stephanie D. Friedman, Adam P. Arkin, Adam M. Deutschbauer, Simon Roux, Vivek K. Mutalik

## Abstract

Single-stranded RNA (ssRNA) and single-stranded DNA (ssDNA) bacteriophages form a key component of the global virome, yet the host genetic networks supporting their infection cycles remain poorly understood. Here we present a genome-wide map of the host factors involved in infection by F pilus-dependent ssRNA and ssDNA phages in *Escherichia coli*. Screens across ssRNA phages spanning all four genogroups of the *Leviviricetes* reveal a conserved network of host dependencies, with the notable exception of the F plasmid gene *traD*. Primary structural receptor components and DsbA-mediated disulfide bond formation are universally required to ensure F pilus integrity, whereas *traD* shows a strict genogroup-specific requirement during entry despite all lineages sharing the same receptor. Gene dosage screens identify elevated copy number of the genomic fragments encoding the HslU protease or the RNA chaperone StpA as barriers that restrict infection. Parallel assays with filamentous ssDNA phages confirm reliance on the host TolQRA complex for entry downstream of pilus engagement and track negative fitness signatures across envelope homeostasis clusters, reflecting the physiological burden of non-lytic virion extrusion. Comparative genomics and high-throughput genetics trace the selectivity of the isolation host, *E. coli* HSF, to a horizontally acquired *Klebsiella* capsule that excludes a large panel of double-stranded DNA phages isolated on other *E. coli* strains while allowing ssRNA and ssDNA virions to engage the extended F pilus and bypass the barrier through native pilus retraction. The work delivers a systematic, class-wide map of single-stranded phage-host interactions, a host platform for accessing uncultured viral diversity, and a functional blueprint for diagnostics, protein antibiotics and biocontrol tools that halt horizontal gene transfer.

## Introduction

Bacteriophages are the most abundant biological entities on Earth and act as critical drivers of microbial ecosystems by shaping host physiology, population dynamics, and horizontal gene transfer^1^. Despite the vast viral diversity uncovered by metagenomic and metatranscriptomic sequencing, classical cultivation frameworks remain heavily biased toward double-stranded DNA (dsDNA) viruses^2^. This historical skew leaves single-stranded RNA (ssRNA) and single-stranded DNA (ssDNA) phages vastly underrepresented in laboratory collections^3^. Recent environmental sequencing efforts show that these small-genome viruses can be highly abundant and active across diverse ecosystems, underscoring that legacy dsDNA model systems provide an incomplete view of phage-host biology^4–10^.

The canonical ssRNA phages, including MS2, Qβ, fr, and R17, were isolated over half a century ago and served as foundational models to solve the mechanics of translation control, virus assembly, and RNA replication^11^. Because these viruses possess minimal coding capacities, they rely entirely on hijacking or manipulating host housekeeping pathways. The cultivated members of this group belong to the *Leviviricetes* class and are classified into four distinct genogroups divided between two genera^12,13^. Strains within a specific genogroup share high (≥80%) nucleotide sequence conservation, whereas sequence identities drop below 40% for strains between genogroups, reflecting extensive evolutionary divergence. Despite their historical utility in molecular biology and their emerging applications in targeted therapeutics and vaccines, no generalizable framework exists to capture or systematically profile the host factor requirements of this divergent viral class across different environmental hosts^4^.

The vast majority of known ssRNA phages specifically depend on the F plasmid-encoded conjugative pilus, a rigid macromolecular filament normally utilized by the host cell for genetic exchange via conjugation^11,14,15^. Decades of research have detailed the precise molecular architecture of this conjugative apparatus, defining the discrete assembly stages, required pathways, and essential genes of a highly dynamic type IV secretion system optimized for both macromolecular transport and extracellular pilus biogenesis^14,16–22,23^. The infection cycle of these F plasmid-dependent RNA phages begins when the viral maturation protein binds tightly to the lattice sides of the F pilus^12,24^ (Fig. 1). Recent structural studies of the MS2-pilus complex demonstrate that this lateral binding event triggers critical conformational modifications that prime the capsid for genomic RNA release^12,25,26^. Following engagement, depolymerization and retraction of the F pilus draw the intact virion toward the cell envelope, initiating a controlled extracellular uncoating process that leaves an empty, intact capsid outside the host^24,27^. This transient exposure renders the viral genome exceptionally vulnerable, giving rise to the characteristic sensitivity of ssRNA phages to external RNase treatment during early entry^11,28^. To protect the genome during translocation across the envelope, the viral maturation protein accompanies the naked RNA into the cell, shielding defined segments from enzymatic degradation^24,25^. However, the exact molecular mechanisms by which this naked RNA ultimately crosses the host cell membranes into the cytoplasm remain uncharacterized.

**Figure 1.**
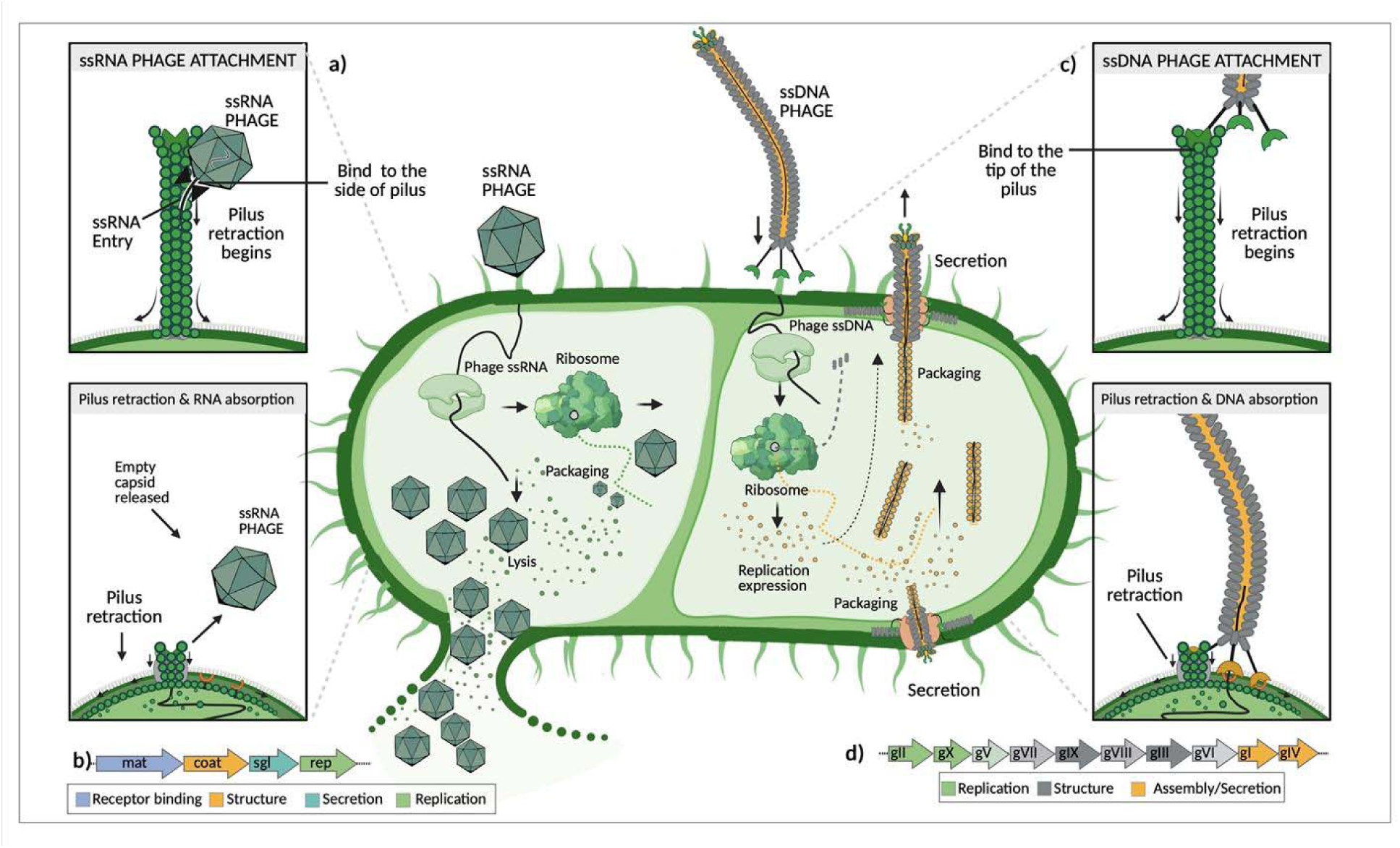
Comparative overview of ssRNA and ssDNA phage infection pathways and genome architectures. (a) Schematic representation of the ssRNA phage life cycle. The ssRNA virion binds to the lateral surface of the retractile F-pilus via the maturation (Mat) protein. Subsequent pilus retraction draws the phage to the host cell surface, initiating the eclipse phase where the capsid is disassembled and the Mat-RNA complex is translocated into the cytoplasm. Following translation and RNA genome replication, mature virions are assembled and packaged until a dedicated lysis protein triggers host cell lysis to release viral progeny. (b) Genomic organization of ssRNA phages. The ssRNA viral genome is encoded within a four-gene architecture: *mat* (codes maturation protein involved in pilus binding), *coat* (codes capsid structural protein), *sgl* (codes lysis protein, Sgl), and *rep* (codes RNA dependent RNA polymerase, RdRP). (c) Schematic representation of the filamentous ssDNA phage life cycle. The filamentous virion initiates infection by binding to the distal tip of the F-pilus via the N2 domain of minor coat protein pIII. Upon pilus retraction, the pIII/pVI terminus of the virion is pulled into the periplasm, facilitating an interaction between the N1 domain of pIII and the host TolA co-receptor. This structural rearrangement drives the injection of the ssDNA genome into the cytoplasm for subsequent rolling-circle replication and phage protein expression without destroying the host envelope. (d) Genomic organization of filamentous ssDNA phages. The ssDNA genome features three functionally segregated gene modules: replication (*gII*, *gX*, and *gV*), virion structural components (*gVII, gIX*, *gVIII*, *gIII,* and *gVI)*, and assembly/secretion machinery (*gI* and *gIV*).

This pilus-dependent entry paradigm stands in stark contrast to the infection cycle of F pilus-dependent, ssDNA filamentous phages, collectively known as the Ff group, which includes M13, fd, and f1^29^. Rather than associating laterally with the pilus lattice, Ff phages utilize their minor coat protein, pIII, to attach exclusively to the tip of the extended F-pilus^30^ (Fig. 1). Retraction of the pilus brings the intact virion into the periplasm, where pIII undergoes structural transitions to directly engage the host inner-membrane TolQRA macromolecular complex^31^. Instead of shedding an empty capsid into the extracellular space, the filamentous viral coat is completely disassembled and recycled into the host inner membrane as the ssDNA genome translocates into the cytoplasm^32^. Furthermore, Ff phages establish a chronic, non-destructive relationship with their host, using rolling-circle replication machinery and cytoplasmic chaperones to continuously assemble and extrude new progeny via specialized membrane secretins without inducing cell lysis^33^.

While there have been some efforts in developing innovative cultivation hosts and indicator strains to improve targeted environmental isolation of plasmid-dependent phages^34,35^, systematic and scalable genome-wide screens that can uncover essential host dependencies and genogroup-specific variations are needed^4^. For example, most historical genetic studies were mapped using only a few Group I phages like MS2 with occasional Group III phage Qβ comparisons^11^. This narrow focus leaves open questions regarding whether F-plasmid host factor requirements apply uniformly across all four genogroups or if there are genogroup-specific dependencies.

Here, we address these open questions by performing a systematic genetic analysis of single-stranded phage-host interactions (Fig. 2a). We assembled a phylogenetically diverse panel of 46 ssRNA phages spanning the known breadth of *E. coli-*infecting *Leviviricetes*. Using parallel random barcode transposon-site sequencing (RB-TnSeq)^36^ and dual barcoded shotgun expression library sequencing (Dub-seq)^37^ technologies, we identified both universally conserved and lineage-specific host factors essential for infection. By deploying F pilus-dependent ssDNA filamentous phages as a concurrent genetic comparator, we mapped key mechanistic divergences between these structurally distinct viral classes as they engage shared host machinery. Our comparative screens validated known structural entry requirements, identified a novel envelope stress regulatory layer mediated by the alternative sigma factor RpoE, and uncovered the capsule-mediated exclusion dynamics that govern the selectivity of the phage isolation platform. Ultimately, this work establishes a comprehensive genetic blueprint for single-stranded phage infection and expands the repository of available isolates, offering a foundational resource for anti-viral defense research and rational design principles to expand viral isolation frameworks.

**Figure 2.**
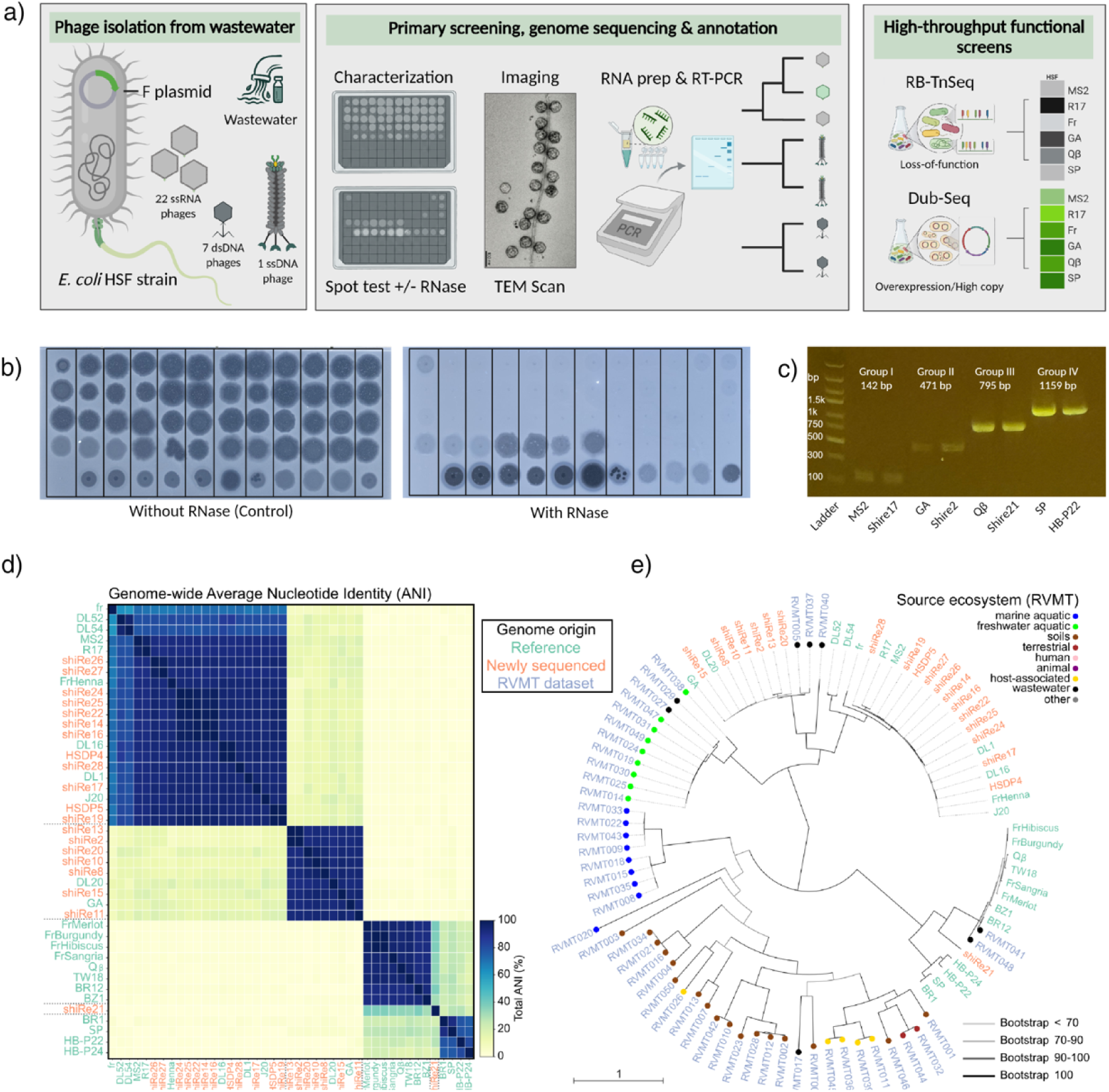
Systematic workflow and experimental pipeline for the isolation, classification, and functional genomics of single-stranded phages. (a) Overview of the phage characterization pipeline. Environmental isolation utilizing the *E. coli* HSF host strain was paired with a compiled library of previosuly isolated phages. Downstream characterization integrated transmission electron microscopy (TEM) structural imaging, RNase A sensitivity profiling, viral RNA extraction, and RT-PCR screening for gel-electrophoretic band size verification. Following whole-genome sequencing, functional profiling was conducted using loss-of-function RB-TnSeq and overexpression Dub-seq genetic screens to map critical host factors driving the infection pathway. (b) Differential plaque assays for plaque-purified candidates. Spot tests performed on the wild-type (WT) *E. coli* HSF host in the presence (+) or absence (−) of 100 µg/ml RNase A. Clearing zones that persist under RNase A treatment signify candidates belonging to either the filamentous ssDNA or double-stranded DNA (dsDNA) phage families. (c) Genogroup classification via RT-PCR. Representative gel electrophoresis of RT-PCR amplification products used to stratify isolated ssRNA phages into distinct, established genogroups based on diagnostic amplicon sizes (Methods, Supplementary Fig. S1). (d) Genome-wide comparison across the ssRNA phage collection. The heatmap displays the genome-wide Average Nucleotide Identity (ANI) across the newly isolated (orange) and references (green) ssRNA viral genomes (Methods). (e) Phylogenetic diversity of the ssRNA phage library and relevant environmental sequences. An RdRP phylogeny was built based on the newly isolated and reference sequences, supplemented by the most closely related metatranscriptome-derived RdRP sequences from the RVMT^6^ dataset (blue labels, see Methods). The ecosystems of origin of RVMT sequences are indicated with colored circles. Branch support values are indicated via the branch color.

## Results and Discussion

### Assembly of a phylogenetically diverse ssRNA phage collection

A major challenge in ssRNA phage isolation is the overwhelming abundance of dsDNA phages in environmental samples, which typically outnumber ssRNA phages and rapidly dominate enrichments on standard *E. coli* strains^2^. To circumvent this limitation, we utilized *E. coli* HS(pFamp)R (hereafter referred to as *E. coli* HSF), a standard indicator strain for environmental monitoring^13,38,39^ that exhibits natural resistance to a broad range of dsDNA phages while maintaining full susceptibility to ssRNA phages due to a specific capsule phenotype characterized below. This selective susceptibility profile enabled the targeted enrichment and isolation of 21 new ssRNA phages from diverse environmental sources, which were complemented by historical as well as recent isolates^34,40^ to yield a final panel of 46 phages (Methods). All isolates were confirmed as genuine ssRNA phages via clear loss of infectivity during exogenous RNase sensitivity testing (Fig. 2, Methods).

RT-PCR genotyping, whole-genome sequencing, and phylogenetic analysis revealed that our compiled collection successfully spans all four established genogroups of F pilus-dependent leviviruses (Fig 2c, Supplementary Fig. S1). All isolates displayed the characteristic compact genome architecture of this class, possessing genomes between 3.5 and 4.3 kb that encode a maturation protein, coat protein, replicase, and for MS2-like genome a lysis protein variant (Fig. 1, Methods, Supplementary Fig. S2, Supplementary Table S1-S2). Transmission electron microscopy of representative isolates from each lineage confirmed the expected icosahedral capsid morphology with uniform diameters of approximately 26 nm (Supplementary Fig. S3). Phylogenetic analysis of the core replicase amino acid sequences indicated that Group I (MS2-like) phages comprised 49% of the collection, followed by Group II (GA-like) at 21%, Group III (Qβ-like) at 21%, and Group IV (SP-like) at 9%, aligning with our RT-PCR genotyping assays using genogroup-specific primers^41^ (Fig. 2d, e). This phylogenetically comprehensive collection (Supplementary Fig. S4) provided a robust experimental foundation to systematically investigate host factor requirements across the *Leviviricetes*.

### Genome-wide genetic screen identifies host factors required for ssRNA phage infection

Having established a phylogenetically diverse *E. coli* ssRNA phage collection, we next sought to comprehensively map the host factors required for infection. To achieve this, we constructed a fully-saturated RB-TnSeq library in *E. coli* HSF strain comprising 208,678 uniquely barcoded transposon insertion mutants (Methods). Deep sequencing of genomic DNA from the unselected baseline library confirmed that insertions were densely distributed across both the host chromosome, with 189,443 insertions mapping to 3,847 genes, and the F-plasmid, with 19,235 insertions mapping to 87 genes in the transfer (*tra*) region. This distribution provided saturating genomic coverage for robust, high-resolution functional genetic analysis (Supplementary Table S3).

Using this barcode-mapped library, we performed genome-wide loss-of-function selection assays across our entire phage panel following our established protocol^42,43^ (Fig. 2a, Methods). Briefly, frozen library aliquots were recovered to mid-log phase and subsequently challenged with individual phages at multiplicities of infection (MOI) of 1 or 10. Cultures were incubated under continuous agitation for eight hours to permit multiple cycles of viral infection, amplification, and stringent selection for phage-resistant mutants. Surviving cells were collected, genomic DNA was extracted from both the pre-infection and post-selection populations, and transposon-adjacent barcodes were amplified for quantification via deep sequencing. Strain fitness scores were calculated as the log2 ratio of barcode abundance at the endpoint relative to the baseline, which were subsequently aggregated to compute gene-level fitness scores^36,44^. In total, we executed 150 independent RB-TnSeq experiments at different MOIs. After cross-referencing with no-phage control experiments to account for baseline fitness effects (Fig. 3, Supplementary Table S3), the fitness scores highlight genes whose disruption confers robust phage resistance reproducibly across biological replicates.

**Figure 3.**
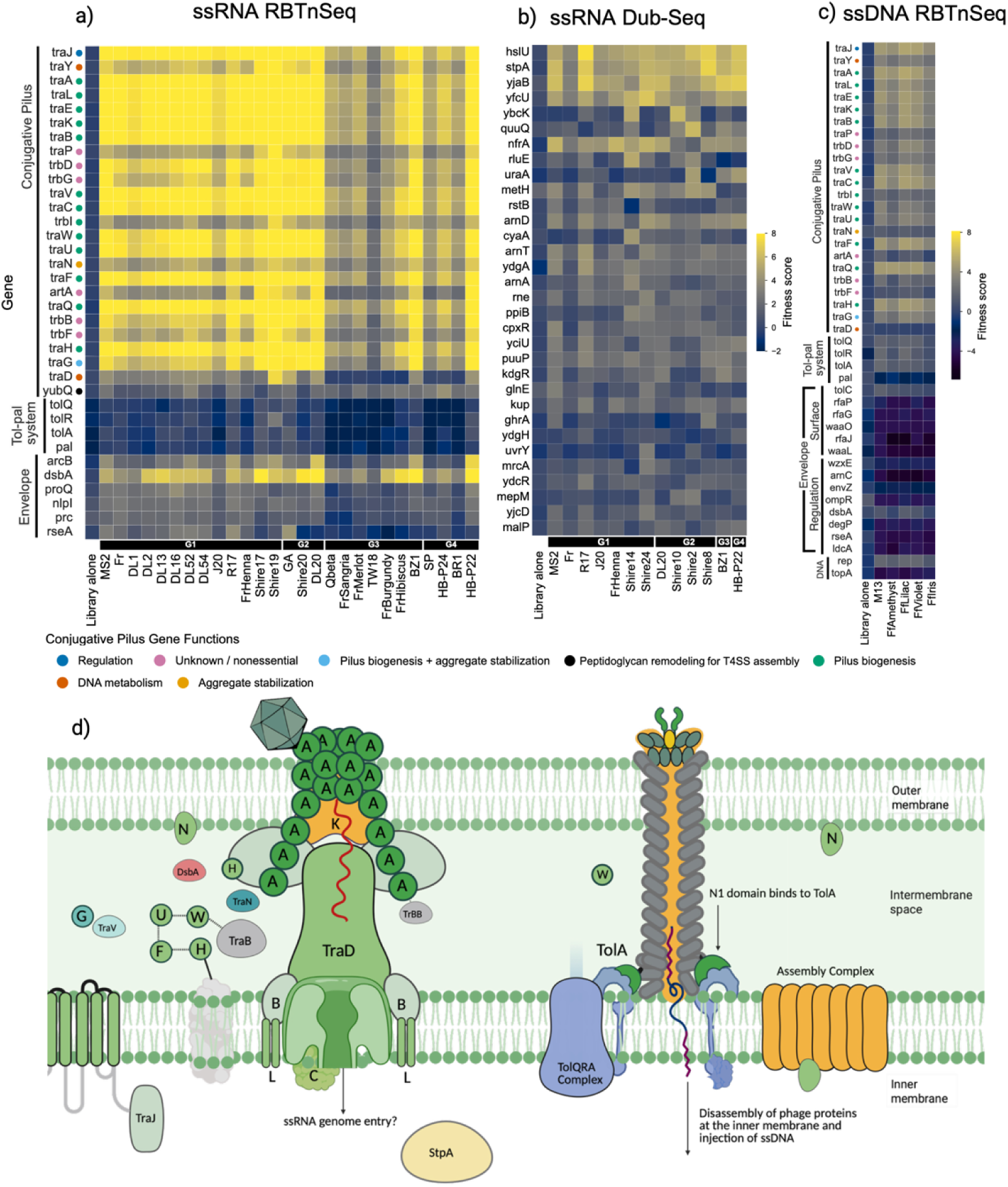
High-throughput functional genomics mapping of host factors required for single-stranded RNA and DNA phage infection. (a) RB-TnSeq fitness profiles of *E. coli* HSF during ssRNA phage infection. Heatmap displaying high-confidence host gene disruption hits across diverse ssRNA phage assays performed at MOI ranging from 1 to 10. Assayed phages are categorized by their respective genogroups, and core host gene functions are annotated on the margin. Top genetic hits required for infection are filtered based on a threshold score achieved in at least one phage assay (Methods). (b) Dub-seq overexpression profiles during ssRNA phage infection. Heatmap showing fitness effects across the representative ssRNA phages and MOI ranges, utilizing the *E. coli* Dub-seq library to identify host factors whose higher copy or expression alters viral propagation kinetics. (c) RB-TnSeq fitness profiles during filamentous ssDNA phage infection. Heatmap illustrating high-confidence host gene disruption hits required for or restrictive to filamentous ssDNA phage infection. (d) Schematic spatial model of the single-stranded phage entry apparatus. Structural diagram mapping the F-pilus assembly, *tra* gene complex products, and auxiliary envelope factors identified in this study across the inner membrane, periplasm, and outer membrane boundaries.

### Systematic profiling identifies genetic determinants involved in ssRNA phage infection pathway

Quantitative filtering of our genome-wide dataset provided a set of functional determinants across the host genome and F plasmid. These high-confidence hits group into a universally required structural apparatus and a discrete cohort of moderate, context-dependent regulatory and protein quality-control factors.

The F plasmid transfer apparatus is required universally across ssRNA phage diversity. Our genome-wide data is in agreement with prior reported observations and shows that the F-type type IV secretion system is required as an integrated, cooperative macro-assembly^12,14–16,22,24,26^. Insertion mutants of genes encoding the physical architecture of the F pilus displayed universally high positive fitness scores across all 27 ssRNA phages tested. Structural subunits and core biogenesis factors, including *traA*, *traL*, *traE*, *traK*, *traB*, *traV*, and *traC*, ranked as top-tier essential hits in every selective enrichment, showing mean fitness scores between 8 and 12 across all viral lineages (Fig. 3a). This profile confirms that functional F-pilus presentation is an absolute, non-redundant prerequisite for infection across all *E. coli* leviviruses tested.

Beyond the physical pilus core, our screens also mapped a stringent requirement for upstream regulatory and assembly factors. The master transcriptional activator *traJ*, which de-silences the 32-kb polycistronic transfer operon by counteracting host H-NS repression at the P_Y_ promoter^45,46^, emerged as a high-scoring hit despite lacking a structural role. Disruption of *traJ* fully abolishes downstream transfer region transcription, mimicking a total structural deletion. The accessory factor *traY*, which modulates P_Y_ output and coordinates the relaxosome complex at the origin of transfer, showed a similarly indispensable phenotype. The dedicated propilin chaperone *traQ* also proved strictly essential, confirming its role in processing propilin prior to filament assembly. Additionally, the specialized periplasmic pilus-extension factors *traF*, *traH*, *traW*, and *traU*, alongside the outer-membrane stabilization genes *traN* and *traG*, were mandatory for infection across all genogroups to convert transient receptor contacts into stable assemblies.

Because the F-plasmid transfer region is transcribed as an extensively coupled polycistronic operon^22,47,48^, internal rho-dependent transcriptional polarity is an inherent regulatory feature of the apparatus that naturally shapes transposon insertion profiles. We observe clear enrichment across uncharacterized loci such as *trbD* and *trbG*, which lack independent downstream promoters, indicating that their genetic disruptions disrupt the structural integrity or stoichiometric balance of the broader macro-assembly.Conversely, *trbF* is structurally insulated from these polar effects due to its independent internal promoter, P*_trbF_*, validating its specific involvement in the ssRNA phage. Rather than evaluating these dependencies as isolated, individual genetic components, our dataset provides a high-resolution, class-wide map of the collective genomic architecture that governs viral entry without requiring exhaustive gene-by-gene complementation. To confirm the essential role of the mature pilin structure and the broader transfer apparatus across all genogroups, we performed targeted, clean genetic validations using an isogenic deletion of the major pilin subunit alongside a total F-plasmid deletion strain (Methods, Fig. 4a, Supplementary Fig. S5), confirming complete resistance to ssRNA phage challenge upon removal of these essential primary docks.

**Figure 4.**
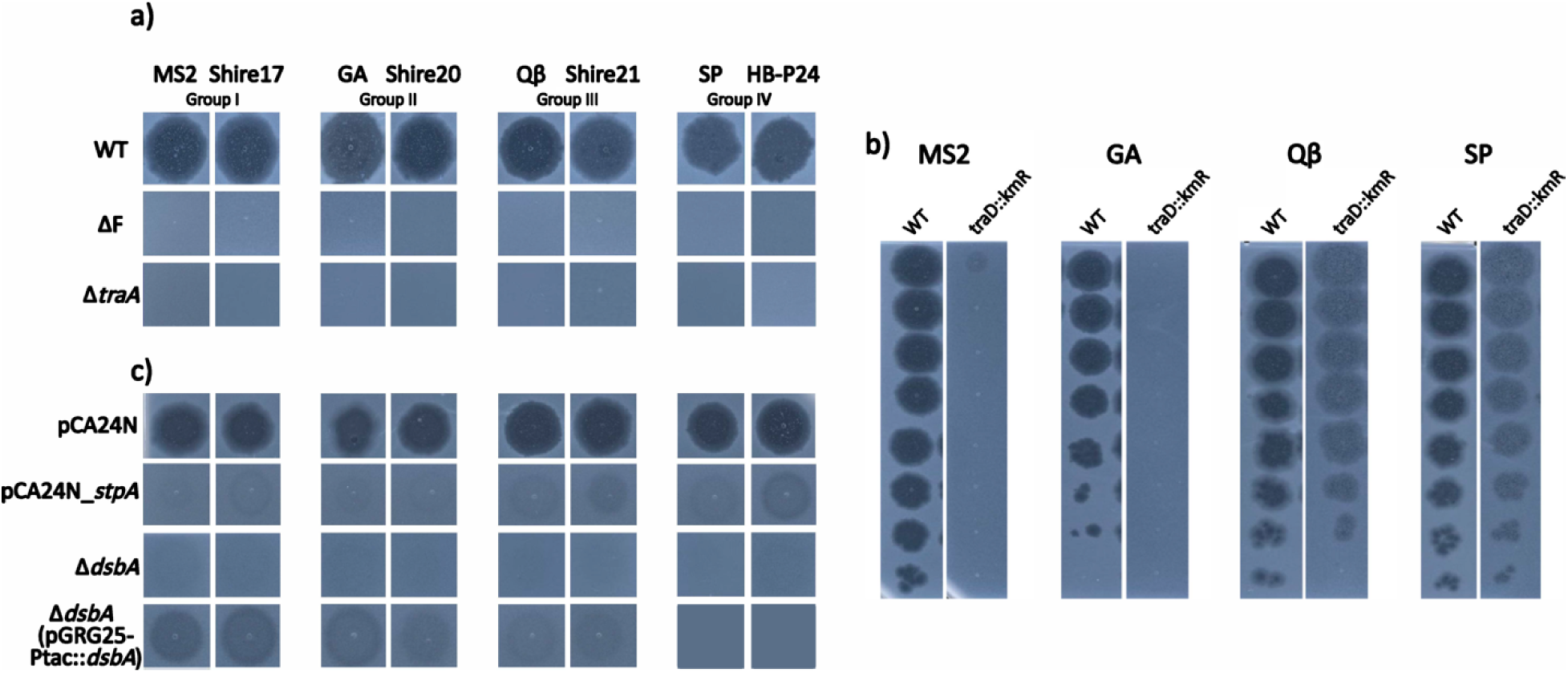
Genetic validation of core host factors and entry machinery across ssRNA and ssDNA phage groups. (a) Comparative spot assays across receptor and assembly mutant strains. Representative plaque clearing assays for phage genogroups 1 through 4 spotted onto lawns of wild-type (WT) *E. coli* HSF and its corresponding genetic deletion backgrounds: F plasmid cured, *traA* deletion mutant (b) EOP assay on the *traD* disruption mutant for all four ssRNA phage genogroups relative to the WT strain. Extended validation profiles and additional plate images are provided in Supplementary Figure S4 (c) Comparative spot assays for phage genogroups 1 through 4 spotted onto lawns of WT *E. coli* HSF with empty pCA24N plasmid, with *stpA* overexpressed, *dsbA* deletion mutant, and complemented dsbA deletion mutant.

### TraD mediates genogroup-specific inner membrane dependency

In contrast to the universal necessity of the pilus filament, the F plasmid gene *traD* exhibited a genogroup-specific fitness profile despite all tested phages utilizing the identical lateral F pilus lattice for initial attachment (Fig. 3a). TraD operates as a type IV coupling protein, serving as a substrate receptor at the cytoplasmic face of the inner-membrane secretion channel. Disruption of *traD* conferred high-confidence phage resistance exclusively to Group I phages in agreement with prior literature^14,49^. Conversely, Group III phages did not show evidence of requirement for *traD*, confirming that Group III viruses can likely bypass this factor entirely during cytoplasmic entry^14,18,24,27^. Our parallel pooling screens revealed that Genogroup II and Genogroup IV phages displayed intermediate *traD* requirements, yielding variable fitness values that failed to establish a concrete baseline requirement. This partial requirement implied that different ssRNA genogroups might exhibit distinct inner membrane translocation dynamics or rely on alternative inner-membrane protein during cytoplasmic genome entry despite sharing an identical primary receptor.

To resolve these screening ambiguities and validate these varying operational requirements across the entire phylogenetic breadth of our collection, we performed efficiency of plating assays with representative phages from all four genogroups against an isogenic *traD* deletion host background (Fig. 4b, Supplementary Fig. S6, Methods). These targeted validation assays clarified the medium fitness scores in our screening noise and recapitulated our high throughput screening metrics. The validation assays demonstrated a strict TraD dependence for Genogroup I and II isolates, whereas Genogroup III and IV phages exhibited unaffected baseline plating efficiencies, revealing that this entry requirement fundamentally trends along distinct viral genera (*Emesvirus* versus *Qubevirus* lineages). While legacy models using temperature sensitive mutants suggested that TraD alters membrane characteristics to stabilize labile Genogroup I replicases rather than participating in literal genome translocation^50,51^, our class wide profiling suggests this differential inner-membrane translocation requirement as a fundamental evolutionary dividing line between the two groups of *E. coli* leviviruses.

### Periplasmic disulfide bond formation is essential for ssRNA phage infection

Among chromosomally encoded host factors, we identified a near-universal requirement for genes involved in the periplasmic oxidative folding pathway. The host gene *dsbA*, which encodes the primary periplasmic thiol-oxidoreductase responsible for catalyzing disulfide bond formation^52–54^, showed strong positive fitness scores across all ssRNA phages tested (Fig. 3a). Operating in tandem with *dsbA*, the plasmid-encoded gene *trbB*, a plasmid-encoded periplasmic disulfide isomerase with DsbC-like activity known to maintain required disulfide configurations of cysteine-rich transfer proteins^55–57^, also scored as a significant hit. These results indicate that proper disulfide bond formation is critical to support ssRNA phage infection.

To validate this functional requirement, we constructed an isogenic *dsbA* deletion mutant and evaluated its susceptibility against a representative panel of ssRNA phages spanning four genogroups (Fig. 4a, Supplementary Fig. S5). The plating efficiency of all tested viral strains on the *dsbA* deleted host was reduced by a thousand-fold to a million fold relative to the wild type strain, confirming that DsbA function is essential for productive infection. We next used geraniol and 2-phenylthiophene to determine if chemical inhibition of DsbA could phenocopy this genetic loss (Methods). Geraniol acts as a competitive inhibitor that directly blocks the DsbA catalytic active site^58^, whereas 2-phenylthiophene derivatives bind the adjacent protein interaction groove to prevent substrate processing^59^. Both chemical interventions restricted phage replication to the same degree as the genetic knockout (Supplementary Fig. S7 and S8), demonstrating that active periplasmic disulfide bond catalysis is required for viral entry.

Host encoded DsbA plays a crucial role in conjugative pilus formation because multiple plasmid encoded transfer factors require proper disulfide bond coordination to fold and stabilize the type IV secretion machinery^56,60^. The F plasmid encoded type IV secretion system proteins are remarkable for their high cysteine content, as TraN, TraU, and TraH contain 22, 10, and 6 cysteines, respectively^57^. For instance, the major pilin subunit TraA requires a disulfide bond between two conserved cysteine residues for structural stability, and assembly proteins such as TraV and TraC similarly rely on oxidative folding. Despite this known requirement for pilus structural biogenesis, a direct functional connection between host DsbA and ssRNA phage infection has not been previously reported. While certain leviviruses (such as Qβ) utilize structural inter-subunit capsid disulfide bonds to enhance virion stability, whereas others (such as MS2) do not^11,61,62^, our screening data revealed a universal block to plaque formation across all assayed genogroups, indicating that host DsbA-mediated pilus assembly remains the dominant functional constraint.

We hypothesize that DsbA acts directly on these multiple cysteine-rich F pilus assembly components^14,20,56^ to ensure the folding of a functional stable macro-assembly capable of binding and translocating ssRNA genomes. Alternatively, beyond its structural role in receptor biogenesis, DsbA may manipulate chromosomally encoded periplasmic factors^63^ during viral genome translocation into the cytoplasm immediately following pilus retraction. Mapping this essential reliance across diverse lineages reveals that ssRNA viral entry is highly sensitive to the native host redox machinery governing host cell envelope structures. This baseline sensitivity underscores how natural variations in the capacity and mechanisms for disulfide bond formation across different bacterial species could shape their susceptibility to viral pathogens.

A secondary class of chromosomal genes, including *nlpI* (outer-membrane lipoprotein adaptor), *prc* (tail-specific protease), *arcB (s*ensor kinase of the ArcAB two-component system), and *proQ* (RNA chaperone), exhibited minor fitness effects across our ssRNA phage pooled fitness assays (Fig. 3a). While several of these host factors, most notably ArcB, are involved in the regulation of F-pilus biogenesis^64–66^ and have been shown to influence MS2 plaque characteristics by modulating receptor abundance^67^, their requirement has not been systematically characterized across different ssRNA phage genogroups. Because our screening model correctly favors essential structural machinery and its core regulators, which drive the strongest phenotypic scores for direct F-pilus components, these secondary envelope stress and auxiliary regulatory factors may not provide a strong enough fitness advantage to be captured in a pooled assay format^42^. Consequently, they do not heavily dictate viral replication dynamics across the lineages evaluated here.

### Genome-scale gene dosage screens identify factors modulating infection efficiency

To complement loss-of-function screens and identify genes whose overexpression or higher copy impacts phage infection, we performed genome-wide gene dosage experiments using Dub-seq library screen (Fig. 1a, Methods). We transferred a previously constructed *E. coli* BW25113 Dub-seq library^37,42^ into *E. coli* HSF and challenged the population with representative ssRNA phages at a MOI of 1 to 10. Cultures were incubated under continuous agitation for eight hours to allow multiple cycles of viral infection and stringent selection. As in the RB-TnSeq assays, we collected survivors, recovered DNA, carried out Bar-seq, and quantified barcode abundance via deep sequencing. Fragment and gene fitness scores were calculated using the log2 ratio of endpoint-to-baseline barcode abundance, as in previous experiments^37,42^.

Our Dub-seq experiments identified a distinct set of loci whose elevated copy number significantly restricted infection efficiency. The top scoring resistance determinants were led by the *hslU* operon, which encodes an ATP-dependent proteasome complex, alongside independent high-fitness scores for the ATPase subunit *hslU* alone (Fig 3b, Supplementary Fig. S9, Supplementary Table S4). In F plasmid biology the HslUV machinery serves as the direct downstream executor of the Cpx envelope stress response pathway by directly interacting with and degrading TraJ which is the master transcriptional activator required for pilus assembly^14,45,68^. Furthermore our results are also in agreement with prior work showing that HslUV acts as a potent multicopy suppressor of multiple single gene lysis (Sgl) proteins including those from F-pilus phages MS2 and non F-pilus phages Hgal1, and PRR1^69^. This indicates a higher copy of *hslUV* acts as a genetic barrier to ssRNA phage infection by two distinct modes. First it represses *tra* operon expression by degrading TraJ and second it provides protection by enhanced degradation of Sgls.

The Dub-seq screens also uncovered a pronounced resistance phenotype for the histone-like nucleoid structuring paralog *stpA*, which protects the host through synergistic transcriptional and post-transcriptional mechanisms (Fig 3b, Supplementary Fig. S9). At elevated copy numbers, StpA is known to form rigid homomeric nucleoprotein bridges or cooperative hetero-filaments with H-NS^70^ to drive robust architectural silencing of the F-plasmid. Due to its high intrinsic affinity for curved, AT-rich DNA tracts, excess StpA can supplement this transcriptional repression to shut down *tra* operon expression and abolish F-pilus receptor assembly^14,45,71–73^. Concurrently, StpA leverages its standalone RNA chaperone domain to facilitate RNA sequence annealing and resolve kinetically trapped tertiary structures ^74^. This post-transcriptional remodeling activity can directly destabilize the conserved viral RNA secondary structures, hairpins, and replicase operator loops needed to coordinate phage translation and replication loops^75^, while simultaneously tuning host regulatory small RNAs to manage the metabolic burden of infection.

Beyond these core proteolytic and nucleic acid architectural networks, the screens revealed a cluster of envelope modification, outer membrane crowding, and inner membrane integrity factors that provide enhanced fitness against ssRNA phages (Fig 3, Supplementary Fig. S9). Because these viruses depend entirely on the host F-pilus type IV secretion system for binding and translocation, higher fitness scores of these surface-altering host factors indicate their role in structural assembly or retraction of the pilus. Specifically, elevated expression of *arnD* and *arnT* alters lipid-A charge to destabilize the anionic lipid microdomains required to anchor the secretion system base^76^, while excess levels of the fimbrial usher protein *yfcU* and the N4 bacteriophage polysaccharide receptor (N4-glycan receptor) secretion protein *nfrA* may physically crowd the outer membrane leaflet to sterically hinder pilus protrusion and viral docking. Similarly, high dosages of the inner membrane protein *ydgA* and the metabolic acyltransferase *yjaB*, alongside the uncharacterized regulatory factor *yciU*, may trigger membrane and metabolic stress that alters the architecture supporting the internal basal rings of the secretion complex.

### Comparative analysis reveals distinct host factor requirements for ssRNA versus ssDNA filamentous phages

The F-pilus serves as the primary receptor not only for ssRNA phages but also for ssDNA filamentous phages of the *Inoviridae* family, including M13, f1, and fd. Although both viral classes strictly depend on the presence of this conjugative appendage, they employ fundamentally different entry mechanisms (Fig. 1). Filamentous phages utilize their minor coat protein pIII to attach exclusively to the distal tip of the F-pilus and subsequently inject their ss DNA genomes through the host TolQRA complex spanning the cell envelope, a pathway that operates independently of pilus retraction^14,29–33^.

To explore how these different entry behaviors translate into specific cellular dependencies, we executed parallel genetic screens using representative filamentous ssDNA phages as a direct control alongside our ssRNA phage panel (Fig 2a, Supplementary Table S1). This comparative analysis uncovered a series of shared and divergent genetic requirements (Fig. 3c, Supplementary Table S5). As expected, both viral types absolutely required the core F-pilus structural genes, such as *traA*, *traL*, *traE*, and *traK*, confirming their mutual dependence on baseline pilus assembly. However, clear genetic divergences emerged immediately downstream of receptor engagement. The F-plasmid gene *traD* yielded universally high fitness scores for Group I ssRNA phages, but was entirely dispensable for ssDNA filamentous infection. This divergence supports the model that TraD regulates entry dynamics essential for ssRNA genome translocation but unavailable for tip-binding filamentous phages. In contrast, components of the host Tol-Pal system, including *tolQ*, *tolR*, and *tolA*, which form the membrane-spanning complex required for filamentous genome translocation, were strictly essential for ssDNA phage infection (Fig. 3c). These same Tol-Pal factors had no detectable impact on ssRNA phage susceptibility (Fig. 3a).

### Parallel screens trace distinct ssDNA filamentous phage requirements and envelope stress dynamics

Beyond validating canonical structural entry complexes, our parallel screens uncovered genetic networks governing host vulnerability during filamentous ssDNA infection (Fig. 3c). The non-lytic nature of the filamentous life cycle alters the selection dynamics relative to traditional lytic selections. Unlike standard lytic phage selections where the rapid amplification of highly resistant mutants creates intense selection pressure that masks broader population depletion trends, the milder physiological burden of continuous filamentous extrusion allowed us to reliably track and interpret high confidence negative fitness scores.

Our screens identified host pathways that protect the cell against the chronic stress of viral propagation, specifically highlighting host envelope homeostasis, membrane modification, and cell-wall maintenance modules. Disruption of the anti-sigma factor gene *rseA* significantly enhanced host susceptibility to ssDNA phage infection, while the gene encoding the periplasmic chaperone *degP*, the *envZ*-*ompR* two component osmolarity sensor system, a specific lipid III flippase, and a murein tetrapeptide carboxypeptidase scored as other high confidence negative fitness hits (Fig. 3c). These principal hits suggest that maintaining tight control over the periplasmic quality control apparatus, lipid homeostasis, membrane architecture, and peptidoglycan cross linking density^77–79^ is important for host cell boundary flexibility during continuous virion extrusion.

Notably, parallel (Dub-seq) gene dosage screening using filamentous ssDNA phages did not yield strong or highly pronounced resistance phenotypes. This lack of a robust over-activation signature likely stems from the non-lytic nature of the filamentous viral life cycle.

### The HSF indicator strain exhibits broad spectrum resistance to classical dsDNA coliphages

Having established the utility of HSF strain for selective viral isolation, we sought to understand the molecular basis for its unusual phage susceptibility profile. The *E. coli* HSF strain has been used as the standard indicator host for FRNA (F-dependent RNA) phage detection in Environmental Protection Agency water quality monitoring protocols for decades^39,40^. A notable observation throughout our isolation efforts was that standard HSF strain cultivation protocols yielded almost exclusively ssRNA and ssDNA phages from environmental wastewater and soil samples.

To systematically test whether this outcome reflects a targeted, broad resistance to dsDNA viruses, we challenged this strain with a laboratory collection of 156 diverse dsDNA somatic phages spanning across at least 6 families, 12 sub-families, and 28 phage genera, all isolated and cultured on diverse *E. coli* strains^43^ (Methods). HSF proved completely resistant to 154 of the 156 dsDNA phages tested (Fig. 5a, Supplementary Fig. S10). This profile stands in contrast to standard laboratory indicator strains such as K-12 MG1655, BL21(DE3), and DH5α, which typically show susceptibility to more than 80% of the phages in this same collection (Fig. 5a). The breadth of this resistance suggested the presence of a robust, generalized physical barrier.

**Figure 5.**
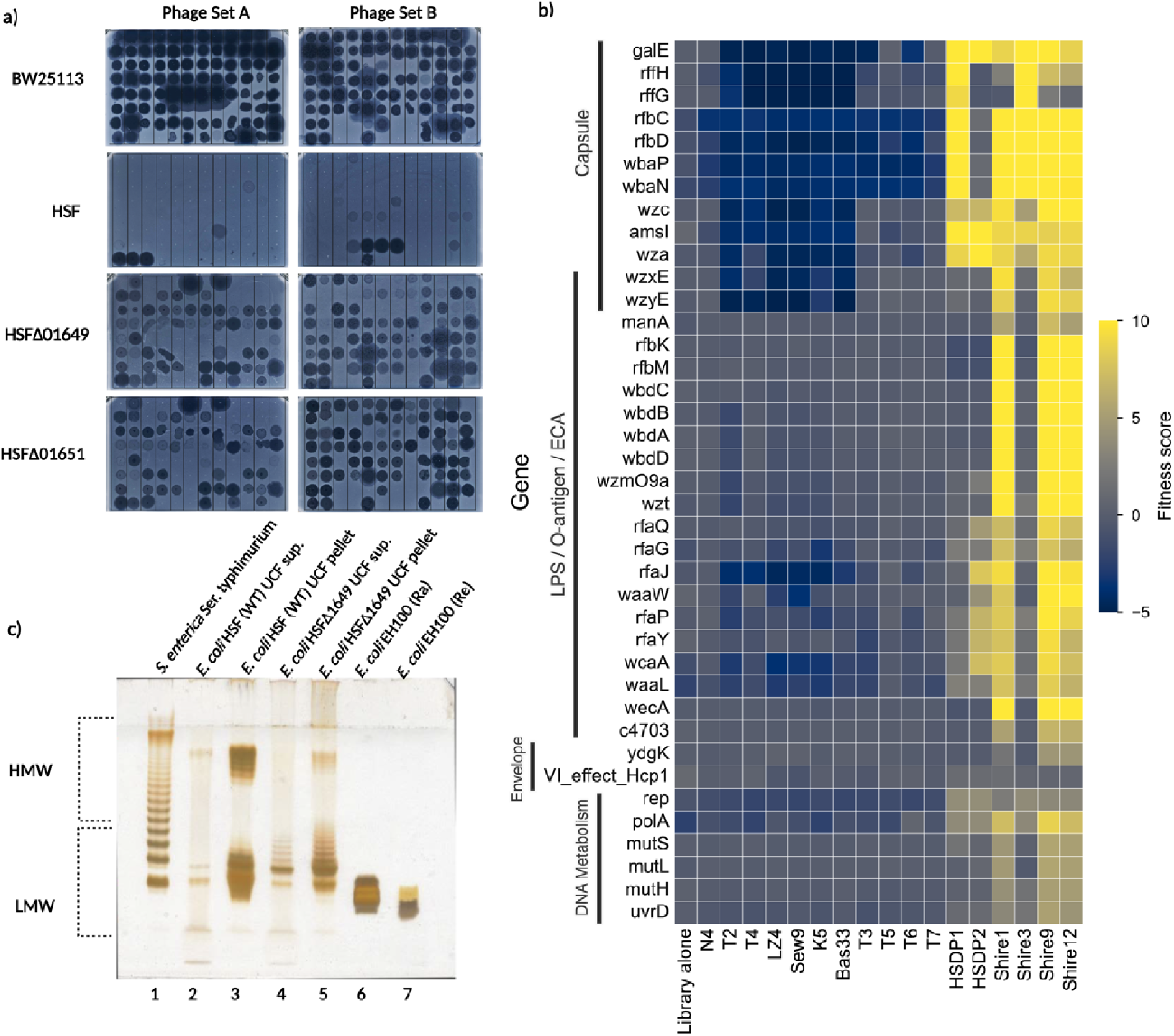
Phenotypic and functional genomic characterization of host envelope features and double-stranded DNA (dsDNA) phage susceptibility profiles in *E. coli* HSF. (a) Comparative dsDNA phage susceptibility profiling and capsule mutant validation. Broad-host-range spot testing comparing the baseline resistance profiles of wild-type (WT) *E. coli* BW25113 and *E. coli* HSF strain background against 156 distinct dsDNA phages partitioned and partially overlapping across Assay Set A (n=93 unique) and Assay Set B (n=66 unique). Exact coordinate layouts of the assayed viral arrays are detailed in Supplementary Figure S10. The bottom panels demonstrate altered validation susceptibility of these 156 phages following deletion of the capsule biosynthetic genes 01649 and 01651. (b) Functional genomic landscape of host determinants required for dsDNA phage infection. Heatmap mapping high-confidence *E. coli* HSF genetic disruption hits derived from RB-TnSeq screens across a diverse collection of dsDNA phages. Screenings were conducted at MOI spanning 1 to 10, with critical host physiological modules and functional categories annotated on the right margin. (c) Deoxycholate-polyacrylamide gel electrophoresis (DOC-PAGE) analysis of co-extracted surface lipopolysaccharide (LPS) and capsular polysaccharide (CPS) isolated from wild-type *E. coli* HSF and the capsule-deficient mutant (Methods). Sample lanes (1 µg total load per lane) are loaded as follows: 1) *Salmonella enterica* subsp. enterica serovar Typhimurium (smooth S-type LPS control); 2) WT HSF post-ultracentrifugation (UCF) supernatant (abbreviated as ‘sup.’); 3) WT HSF post-ultracentrifugation pellet; 4) *E. coli* HSF strain with 01649 deletion post-ultracentrifugation supernatant; 5) *E. coli* HSF strain with 01649 deletion post-ultracentrifugation pellet; 6) *E. coli* EH100 (rough Ra-type LPS mutant control); and 7) *E. coli* EH100 (deep rough Re-type LPS mutant control). Low-molecular-weight (LMW) glycolipid (see lanes 6 and 7). High-molecular-weight (HMW) glycolipid bands are indicated.

### Identification of *Klebsiella*-like capsular locus as a divergent entry determinant

To uncover the molecular basis of this broad physical barrier, we performed a comparative genomic analysis of HSF against standard laboratory *E. coli* strains BW25113 and C-3000, a classic permissive host for ssRNA phages (Methods). The search for candidate resistance genes identified 193 strain-specific genes in the HSF genome, with 95 genes scattered along the chromosome and 98 genes located across seven distinct genomic clusters. While four of these clusters were associated with predicted prophages, two with type III and type VI secretion systems, and one with an integrative conjugation element, the eighth HSF-specific gene cluster (genes MJ392_08245 to MJ392_08320 in CP092639) was discovered directly adjacent to the *galF* gene. In standard *E. coli* K-12 genomes, this locus natively houses the colanic acid biosynthetic operon. However, the HSF genome entirely lacks typical colanic acid machinery, encoding instead a completely divergent 23 kb polysaccharide biosynthetic module situated between the *galF* and *ugd* genes that includes unique glycosyltransferases, polymerases, and export machinery.

A nucleotide sequence similarity search in the NCBI core_nt database^80^ identified this cluster as a homologous match to the capsular polysaccharide locus of *K. pneumoniae* capsular type K47. Pairwise BLASTP analysis demonstrated that seventeen HSF capsular genes share 90% to 99% amino acid identity with the proteins encoded by the *K. pneumoniae* K47 locus (Supplementary Table S6). Only the cpsACP gene of the K47 locus, encoding a putative acid phosphatase, is truncated and rendered non-functional in HSF due to a transposase insertion. This specific *Klebsiella*-like capsular operon is exceedingly rare within the broader *E. coli* lineage, appearing in only 115 out of 65,704 available *E. coli* genomes in the NCBI RefSeq database (as of August 2026), compared to 528 genomes of *Klebsiella* spp. Architecturally, the HSF capsular locus is flanked upstream and downstream by partial IS1 variant transposase sequences. The strategic positioning of these mobile elements strongly implies that this dense carbohydrate synthesis cluster was horizontally introduced into the parental HS lineage via an IS1-mediated insertion mechanism.

To determine if alternative immune networks contributed to this phage resistance phenotype of *E. coli* HSF, we cross-referenced the strain-specific gene list against phage defense predictions generated by DefenseFinder-v2^81^. No strain-specific intracellular defense systems were found in the CP092639 genome. Notably, multiple alignments revealed that our host sequence lacks a substantial 60-kilobase segment containing the bacteriophage exclusion system (BREX)^82^ natively present in the classical HS reference genome (NCBI accession number NC_009800). This missing region, flanked by identical transposase insertion elements, contains the exclusion cluster, one or more prophages, and several accessory loci. This DNA segment is similarly absent from the reference genome of *E. coli* HSF (ATCC 700891) as well as our in-house stocks, validating that our single-strand selective host lacks an operational BREX network. Consequently, the broad-range phage-resistant phenotype of the HSF strain cannot be explained by known intracellular defense networks, pointing instead to the horizontally acquired capsular matrix as the primary candidate.

### Functional mapping of reciprocal surface matrix requirements via high-throughput screens

Given that this unique surface mask closely resembles a *Klebsiella* capsule, we reasoned that the HSF strain might remain vulnerable to specialized environmental phages naturally adapted to recognize or degrade this specific sugar architecture. To test this hypothesis, we performed wastewater enrichments utilizing HSF as the isolation host (Methods). From these enrichment trials, we successfully isolated lytic phages from multiple plaques and resolved their taxonomic identities via whole-genome sequencing (Methods). Genomic analysis revealed that these isolates comprise dsDNA phages sharing high sequence similarity with known *E. coli*-, *Salmonella*-, or *Klebsiella*-infecting lineages (Supplementary Table S1), with their closest sequenced relatives associated with established *Klebsiella* (Shire3, Shire9, and Shire12), *Salmonella* (Shire1), and *Escherichia* (HSDP1 and HSDP2) phages. Crucially, these dsDNA phages were co-isolated from the exact same wastewater samples utilized for our single-stranded (ssRNA and ssDNA) phage discoveries. This confirms that the HSF strain can support active replication of diverse dsDNA viral classes sharing the same ecological niche.

To test how this capsule type physically restricts typical dsDNA coliphages while remaining permissive to these environmental viruses, we performed genome-wide screens with our dsDNA phage collections against the *E. coli* HSF RB-TnSeq library (Fig. 2a). When we assayed 11 diverse *E. coli* dsDNA phages, mutations within the capsular biosynthesis cluster scored significant negative fitness scores^83^, indicating that the native capsule protects the host and that deleting these genes allows classical coliphages to infect (Fig. 5b). In contrast, when we screened the *Klebsiella*-like dsDNA phages isolated from our environmental wastewater enrichments, capsular genes and O-antigen loci scored as strong positive fitness hits (Fig. 5b, Supplementary Table S7). This reciprocal scoring pattern demonstrates that our newly isolated dsDNA phages strictly require the surface capsule and O-antigen matrix for productive infection, as host mutants lacking these structures survive the viral challenge.

This divergent profiling highlighted an interconnected network of capsular polysaccharide, lipopolysaccharide core, and lipid processing factors that are essential for environmental phage infection but protective against classical coliphages. Within the K47 capsular region, key positive fitness hits for the environmental isolates mapped to the complete dTDP rhamnose biosynthetic pathway, regulatory assembly elements such as the capsule tyrosine kinase and phosphatase, and multiple peripheral glycosyltransferases. This surface dependency extended beyond the capsule operon to include essential core lipid and O-antigen processing machinery such as phosphomannomutase, O-antigen ligase, and Lipid III flippase (Fig. 5b). Furthermore, losing outer lipopolysaccharide core modifications via mutations in the *rfa* cluster genes *rfaQ*, *rfaG*, *rfaP*, and *rfaY* conferred absolute resistance to the environmental isolates, confirming that these specialized environmental phages require a fully intact, native cell surface matrix for successful attachment and entry.

### Structural modeling of tail fiber adaptations driving capsule recognition

Because phages HSDP1 and HSDP2 were the only *E. coli* dsDNA phages from our baseline collection able to bypass this barrier and infect the wild-type HSF strain, we sought to investigate the structural determinants underlying their unique host range. Protein-coding genes within the tail loci of both phages were retrieved, aligned with homologous viral sequences, and modeled using AlphaFold3 to map specific functional domains (Methods). Phage HSDP2 belongs to the *Tequatrovirus* genus and is closely related to classical T2 and T4 phages. It possesses a typical T4-like gp37 long tail fiber receptor-binding protein; however, structural modeling revealed a unique 152-amino acid domain located directly upstream of the collar domain that is completely absent from other T4-like phages (Fig. 6A–B). While this domain lacked canonical annotations, a BLASTP search yielded a high-confidence match to a tail fiber protein from a *K. pneumoniae* strain KAB03 genomic scaffold (WP_180106116.1, 86% identity, 100% coverage), suggesting that this domain may harbor acquired depolymerase activity targeted against *Klebsiella* K47-type capsules.

**Figure 6.**
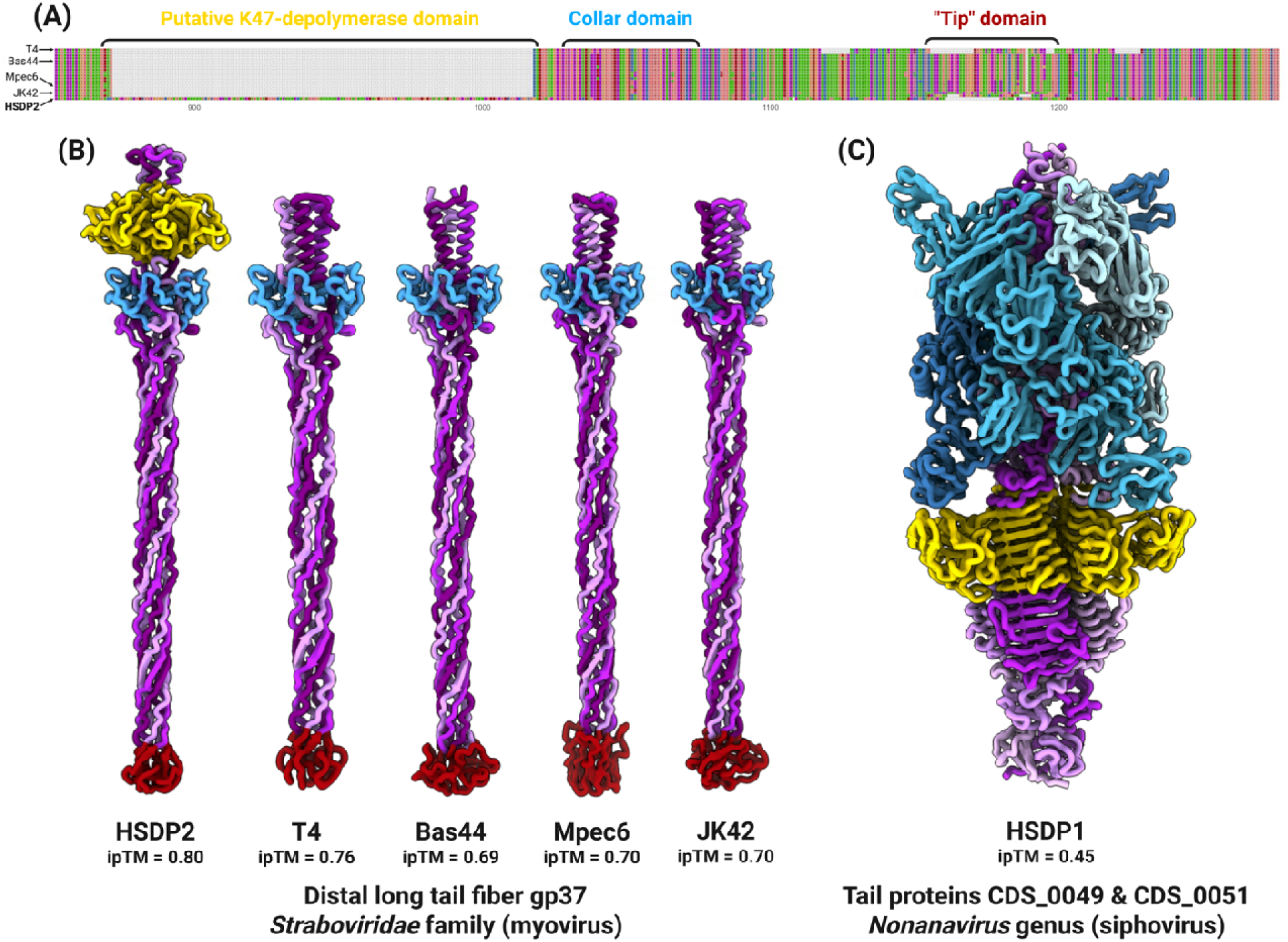
Putative K47-specific depolymerase domains in *E. coli* dsDNA phages HSDP1 and HSDP2. (A) Protein alignment of the C-terminal section of gp37 long tail fiber receptor-binding proteins from T4-like phages. Relevant structural domains are indicated with brackets and colors corresponding to their localization in (B) AlphaFold3 models representative trimeric gp37 C-terminal sections. Purple indicates monomers, the receptor-binding “tip” domain is highlighted in red, the collar domain in blue, and the putative K47-specific depolymerase domain of phage HSDP2 in yellow. (C) AlphaFold3 modeling of phage HSDP1 tail spike complex, made of 2 trimeric tail proteins (PCOOMCAM_CDS_0049, in blue, and PCOOMCAM_CDS_0051, in purple). The homologous K47-specific depolymerase domain is highlighted in yellow.

Conversely, the second permissive dsDNA phage, HSDP1, is a siphovirus belonging to the *Nonanavirus* genus. Protein domain analysis revealed that one of its primary tail proteins displays significant homology (32% identity, 41% coverage) to the tail spike protein Dpo43 from *Klebsiella* phage vB_KpnP_IME205, which exhibits K47-specific capsule depolymerase activity. AlphaFold3 structural modeling of the HSDP1 tail spike shows that this putative capsule depolymerase domain is located directly upstream of the tail tip (Fig. 6C). In both instances, comparative genomics and protein domain analysis allowed us to identify unique putative depolymerase domains, suggesting how these two *E. coli* dsDNA phages bypass the K47 capsular shield of the HSF strain.

### Genetic validation and structural architecture of the capsule entry barrier

We next performed a clean deletion of two key genes (polysaccharide export lipoprotein encoding *wza* (OHPLBJKB_01649) and tyrosine-protein kinase encoding *wzc* (OHPLBJKB_01651)) within the capsular operon to validate these opposite screening phenotypes. The resulting capsule gene deletion strains showed restored susceptibility to the majority of classical dsDNA coliphages, while completely resisting infection by the newly isolated capsule dependent environmental phages (Fig. 5a, Supplementary Fig. S10). This genetic testing confirmed that capsule expression is necessary for the distinct resistance and susceptibility phenotypes. Notably, the capsule deletion strain maintained full susceptibility to ssRNA phages across genogroups (Supplementary Fig. S11), confirming that capsule presence does not affect ssRNA infection kinetics.

To validate the biochemical and structural changes underlying this capsule dependent susceptibility phenotype, we performed electrophoretic and chemical compositional comparison of the isolated cell surface glycolipids from the *E. coli* HSF wild type and the Δ01649 capsular deletion mutant (Fig. 5c, Supplementary Fig. S12, Supplementary Tables S8 and S9). We found that disruption of the capsular locus causes a major loss of a higher molecular weight polysaccharide polymer enriched broad band in PAGE, paired with the appearance of a prominent, lower molecular weight ladder in the HSF Δ01649 mutant (Fig.5c). Composition analysis supported these electrophoretic changes, by demonstrating a significant reduction in overall mannose content from 52% to 20% in the capsule deficient strain accompanied by a simultaneous increase in the relative content of rhamnose, galactose and fatty acids (Supplementary Fig. S12) that could indicate a relative increase in the content of molecules due to the LPS. No major changes were detected in the relative content of fatty acids. Notably, the total carbohydrate content in the isolated glycolipids in the mutant dropped to 43% from the 81% found in the wild-type (Supplementary Tables S8 and S9). Taken together, these data imply that the native capsule consists of a mannose-enriched polymer matrix that normally acts as a binding interface (“mannose shield”) for capsule-dependent phages while masking the underlying rhamnose-enriched structures, likely associated with LPS or other surface glycolipids^84^.

Cryo-electron tomography reconstructions provided a clear visual explanation for this selective entry barrier (Fig. 7a-b). The reconstructions revealed that the long F-pili extend 2 to 5 micrometers from the cell surface, allowing them to penetrate through and reach far beyond the dense capsular matrix which extends only 50 to 150 nanometers from the outer membrane. This distinct structural layout allows small ssRNA and ssDNA virions to easily engage the extended pilus tips and traffic toward the outer membrane via pilus retraction. These results provide a structural explanation for the selective enrichment of single-stranded phages from environmental samples, where the horizontally acquired *Klebsiella* derived capsule selectively shields somatic receptors while the extended F-pilus structure serves as an entry route for pilus dependent viral classes.

**Figure 7.**
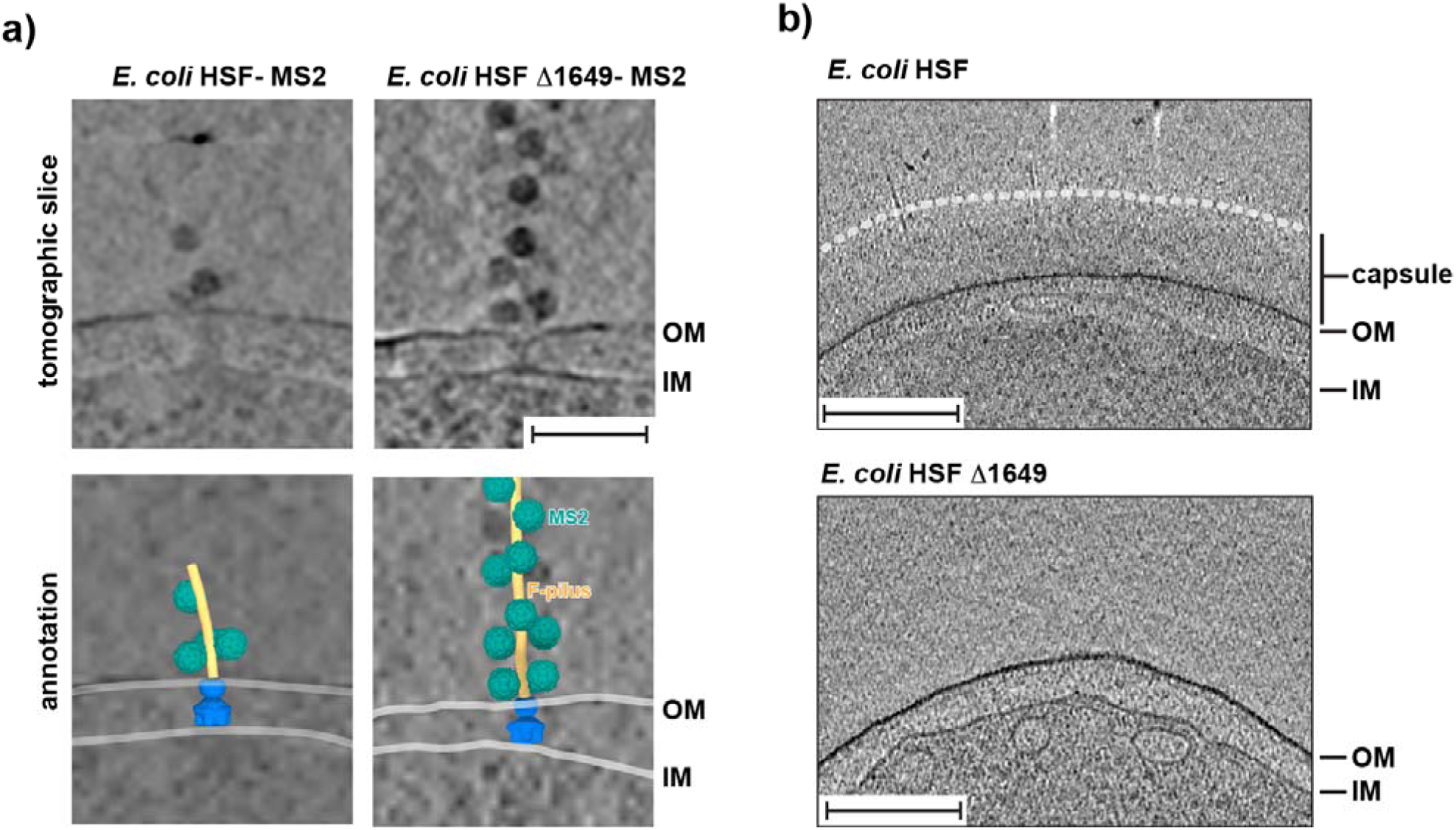
In situ tomographic visualization of ssRNA phage interaction with encapsulated and unencapsulated host envelopes. (a) High-resolution spatial mapping of virion-pilus engagement. Representative electron tomograms (top row) and corresponding three-dimensional segmented annotations (bottom row) displaying ssRNA MS2 phage particles (green) bound to F-pili filaments (yellow). Virion-pilus complexes are captured in close geometric proximity to the host cell envelope across both the wild-type *E. coli* HSF strain and the capsule biosynthetic knockout strain (*E. coli* HSF Δ01649) (b) Phenotypic comparison of capsule architecture. Visual comparison tracking structural envelope features between capsular (top) and non-capsular (bottom) *E. coli* strains. Structural boundaries are labeled as follows: OM, outer membrane; IM, inner membrane. Scale bars represent 100 nm.

## Summary and conclusions

This study provides a comprehensive characterization and genome-wide mapping of host determinants governing ssRNA and ssDNA phage infection. By pairing systematic loss-of-function and overexpression screens, we mapped the precise genetic landscape regulating infection cycle of atypical phages. We established that ssRNA phage infection depends on primary disulfide bond catalysis managed by host *dsbA*, which coordinates the oxidative folding of highly cysteine rich transfer machinery subunits such as TraN, TraU, and TraH to build a stable type IV secretion system pilus. Overexpression screens with ssRNA phages revealed that elevated dosages of the fragment encoding cytoplasmic protease *hslUV* actively restrict infection, whereas ssDNA phages show minimal overexpression resistance probably due to their non-lytic extrusion lifestyle. Furthermore, our comparative framework resolved the molecular basis of the selective isolation profile of the *E. coli* HSF indicator host. This selective barrier is driven by a horizontally acquired *Klebsiella* capsule locus that acts as a physical shield against classical dsDNA coliphages, while simultaneously functioning as an essential entry receptor for adapted environmental *Klebsiella* phages. Electrophoretic, chemical, and cryo-electron tomography analyses confirmed that this high molecular weight, mannose rich polymer matrix effectively blankets somatic receptors, but remains fully permissive to ssDNA and ssRNA viruses that utilize long, protruding F-pili to bypass the shield and traffic to the cell surface via native pilus retraction.

In summary, classical genetics has leveraged single-stranded phage infection for decades as a highly sensitive diagnostic readout for intact, operational conjugation machinery^15^. Our systematic screening framework effectively scales this historical principle into a comprehensive functional mapping platform deployed across diverse phage lineages. By directly pairing these genomic datasets with comparative selections, our screens reveal how distinct viral classes target common host components under completely divergent replication and entry paradigms. As a unique and large-scale isolation and functional mapping of these minor viral classes, this collection provides an extensive biological resource to resolve long standing ecological and mechanistic challenges in studies of atypical viruses. Access to cultivated representatives expands our fundamental understanding of plasmid-phage interaction determinants, enabling systematic gene essentiality studies and the discovery of novel single gene lysis proteins functioning as minimalist protein antibiotics^69,85–87^. Mechanistically, this collection serves as a tractably defined system to dissect how viral genome comprising single-stranded nucleic acids traverse complex lipid and peptidoglycan boundaries during entry and egress steps. Furthermore, pairing these physical isolates with high throughput phenotypic scores establishes a high confidence training dataset for computational machine learning approaches^43,88^.

Beyond expanding fundamental biology, these diverse single-stranded phage model systems provide a versatile toolbox for next generation biotechnological and diagnostic materials^11^. The collection spans across a broad structural palette of unique capsid architectures that can be engineered as modular nanobiotechnology scaffolds or targeted vehicles for payload delivery to specific members of a microbial community. In particular, expanding our catalog of diverse ssRNA phages is important towards development of new RNA binding protein technologies, scaling the utility of canonical model systems like MS2 and PP7 into an array of programmable orthogonal tools for transcript imaging, RNA tracking, and synthetic biology circuits^89,90^. Their robust, simple capsids can also serve as stable biological surrogates to design advanced diagnostic materials, including environmental sensor arrays and protective mask substrates optimized to block airborne viral transmission. Finally, harnessing the absolute reliance of these atypical viral classes on retractile pili offers an elegant functional framework to rationally manipulate microbiomes, establishing a target specific biocontrol platform to selectively eliminate mobile plasmids and halt the horizontal gene transfer of antibiotic resistance genes across environmental reservoirs.

## Materials and Methods

### Bacterial strains and growth conditions

The bacteriophages used in this study and their sources are listed in Supplementary Table S1. Information on all bacterial strains, plasmids and primers used in this study can be found in Supplementary Table S2. All enzymes were obtained from New England Biolabs (NEB), and primers were synthesized by Elim Biopharmaceuticals, Inc. (Hayward, CA). Unless stated otherwise, bacteria were grown in LB medium supplemented with appropriate antibiotics at 37°C with shaking. All bacterial strains were stored at -80°C for long-term storage in 15% sterile glycerol (Sigma). *E. coli* HS(pFamp)R (ATCC 700891) was obtained from ATCC and used as the primary host strain throughout this study. The genetic characteristics of *E. coli* HSF have been described previously^39^. This strain contains three antibiotic resistance markers: ampicillin on the Famp plasmid responsible for pilus production, and streptomycin and nalidixic acid on the chromosome. *E. coli* HSF was cultured in LB Lennox broth (10 g/L tryptone, 5 g/L NaCl, 5 g/L yeast extract) or LB agar (LB Lennox with 1.5% Bacto agar) supplemented with 100 μg/mL nalidixic acid (NAL), 15 μg/mL ampicillin (Amp), and 15 μg/mL streptomycin (Str). *E. coli* strains transformed with plasmids were selected in the presence of 100 μg/mL kanamycin (Kan), 100 μg/mL NAL, 15 μg/mL Amp, 15 μg/mL Str, or 30 μg/mL chloramphenicol (Cam).

For experiments involving phage propagation on agar medium, MS2-specific medium^91^ was used with slight modifications. This medium contained 10 g/L tryptone, 1 g/L yeast extract, and 8 g/L NaCl with 1% Bacto agar for bottom agar and 0.45% Bacto agar for top agar. After autoclaving and cooling to 50°C, the medium was supplemented (‘MS2 media supplements’)^91^ with 0.1% glucose, 2 mM CaCl□, and 10 μg/mL thiamine.

### Isolation and propagation of phages

All phages were isolated and propagated using *E. coli* HSF strain. Due to the high propensity of these single-stranded phages to cause cross-sample contamination, strict containment protocols were enforced; all phage handling and manipulations were performed exclusively within a Class II Biosafety Cabinet and tubes were opened strictly one at a time.

Environmental samples were collected from wastewater, surface waters, and agricultural sources in California. Solid and semi-solid samples were dissolved in LB medium to achieve saturation and filtered through 0.22 μm syringe filters before enrichment. Liquid samples were mixed with an equal volume of 2X LB broth, supplemented with 10 mM MgCl□ and appropriate antibiotics (100 μg/mL NAL, 15 μg/mL Amp, 15 μg/mL Str). Overnight cultures of *E. coli* HSF (100 μL) were added to the mixture and incubated overnight at 37°C with shaking at 50 rpm.

Enriched cultures were centrifuged at 5,000 rpm for 10 min at 4°C, and supernatants were sterilized by filtration through 0.22 μm filters. Phage presence was assayed by double agar overlay method. Briefly, serially diluted enriched samples were mixed with 4 mL of 0.45% soft agar containing MS2 media supplements, 10 mM MgCl□, and antibiotics, then poured over bottom agar to obtain individual plaques. Phages were purified at least twice by picking individual plaques, suspending them in SM buffer (Teknova) supplemented with 10 mM CaCl□ and 10 mM MgSO□ (Sigma), and replating on bacterial lawns. Phage titers were determined by spotting 2 μL of 10-fold serial dilutions in the SM buffer. Purified phages were stored at -80°C in the presence of a bacterial host and 8% DMSO for long-term storage, and as filter-sterilized (0.22 μm) lysates at 4°C for immediate use. Typical stability of these ssRNA phages lasted for 3 months at 4°C.

For initial validation of phage genome type, serial dilutions of each phage were spotted on designated media plates with bacterial lawns containing 100 μg/mL RNase A in the soft agar, with SM buffer alone used as a control. Complete inhibition of plaque formation on RNase-containing plates indicated ssRNA phages. Phage identity was further validated by transmission electron microscopy (TEM) and whole-genome sequencing. For TEM, 10□ µL of lysate was placed on to 200 mesh formvar-covered, carbon-coated copper grids (EMS, Hatfield, PA, USA), incubated for 1□ min, briefly rinsed with ultra-pure water, then stained with 2% uranyl acetate for 2□ min. Samples were imaged on Jeol 1400-FLASH 120KV TEM and a Hitachi HT7800 120 kV TEM. Imaging was performed at a magnification range of 12 to 60kX with the Oneview 16-Megapixel camera (Gatan®).

In order to isolate dsDNA phages HSDP1 and HSDP2 infecting *E. coli* HSF, the standard protocol was modified by adding 100 µg/mL RNase A in the media for phage enrichment and isolation steps. This enzymatic selection process was not applied to the other dsDNA phages (Sheri 1, 3, 9 and 12) isolated during the screens.

### Complete genome sequencing and assembly

Pure phage lysates with titers of at least 10□ PFU/mL were used for nucleic acid extraction. To remove host DNA contamination, phage lysates were incubated with DNase I (NEB) for 30 min at 37°C. RNA was extracted using the MagMAX Viral RNA Isolation Kit (Thermo Fisher Scientific) following the manufacturer’s instructions. RNA concentration was determined using a Qubit 3.0 Fluorometer with the Qubit RNA HS Assay Kit (Thermo Fisher Scientific).

RNA genomes were sent to SeqCenter (PA) or NGSLAB (NY) for reverse transcription and sequencing. Alternatively, cDNA synthesis was performed in the lab following the manufacturer’s recommended protocols. cDNA was synthesized using the Induro® Reverse Transcriptase (NEB) with 300-500 ng of genomic RNA. This reaction mix was then incubated for 2 min at 25°C, followed by 10 min at 55°C. Enzymes were inactivated by heating the reaction mix at 95°C. The cDNA product was stored at -20C.

Sequencing was performed on Illumina NovaSeq or iSeq platforms to produce 150 bp paired-end reads. Sequencing reads were adapter trimmed (NexteraPE adapters) and quality filtered with bbtools v39.52 (tools bbduk.sh and polyfilter.sh, options tbo tpe hdist=2 k=23 mink=9 hdist2=1 minlen=50 ktrim=r, and qtrim=r trimq=20 maq=20). For samples with high read numbers, reads were downsampled to 50,000 reads using seqtk to facilitate assembly. The resulting reads were assembled with SPAdes v.3.15.4 (options --isolate --only-assembler -k 21,33,55,77,99,121). Sequencing reads were deposited in the NCBI Sequence Read Archive (SRA). Accession numbers for previously published genomes and those newly generated in this study (being submitted to NCBI) can be found in Supplementary Table S1.

### Genogroup characterization through reverse-transcription PCR (RT-PCR)

To categorize ssRNA phages based on their genogroup, the Qiagen One-Step RT-PCR kit was utilized with genogroup-specific primers^40,41^. Each of the four primer sets binds to only one of the four ssRNA genogroups, with each product amplifying to a different size to ensure they are easily distinguishable. Group I yields 142 bp, group II yields 471 bp, group III yields 795 bp and group 4 yields 1159 bp^41^. Each ssRNA phage was run against all four primer sets (Supplementary Table S2), with the expectation that amplification would only occur in their respective genogroups.

A 5 µL volume of each phage was aliquoted into a sterile PCR tube and incubated at 98°C for 5 minutes, allowing heat to release the RNA genome. Following heat release and a 2-minute incubation on ice, the following reagents were added to each sample: 10 µL of 5X Qiagen Reaction Buffer (Qiagen), 2 µL of 10mM dNTP (NEB), 0.5 µL of RNase inhibitor (Ambion), 1 µL of 10 µM genogroup-specific FRNA forward primer, 1 µL of 10 µM genogroup-specific FRNA reverse primer, 2 µL of RT-PCR enzyme (Qiagen), and 28.5 µL of nuclease-free H2O. In the thermocycler, samples underwent the reverse transcription step for 30 minutes at 50°C before cycling 40 times between 94°C, 55°C, and 72°C for 1 minute at each temperature.

Following RT-PCR, amplicons were run on gel electrophoresis using an E-Gel PowerSnap (Invitrogen) and a 1% agarose gel. Genogroup-specific size-based separation of amplicons was observed under UV light with an E-Gel DNA Express Ladder (ThermoFisher). As a negative control, a template-less RT-PCR reaction was prepared for each genogroup primer master mix.

### Phylogenetic and comparative genomics analysis

To construct complete genome-based phylogenetic trees, a collection of 23 previously published phage genomes retrieved from NCBI and 20 newly sequenced phages were analyzed. Genome-wide similarity was computed for each pair of genomes as total ANI (“tani”) with vclust v1.3.1^92^. For the phylogenetic tree, relevant sequences from the RVMT dataset^6^ were first recruited based on a BLASTP^93^ (v2.17.0+, default parameters) search of the RdRP protein sequences predicted from our 43 genomes. The top 20 RVMT hits per query were selected, deduplicated, and aligned with the RdRP sequences from our 43 genomes with MAFFT v7.526 (--auto parameter)^94^. The phylogenetic tree was then built from this alignment with IQ-TREE v3.1.1 (-m MFP -bb 1000)^95^. The genomic similarity heatmap was rendered with seaborn v0.13.2^96^, while the phylogeny was visualized with the ete3 toolkit v3.1.3^97^.

### Construction of *E. coli* HSF RB-TnSeq library

The *E. coli* HSF transposon mutant library was constructed via conjugation of *E. coli* HSF with *E. coli* AMD290, which carries the mariner transposon vector library^36,44^. Both strains were grown overnight in LB at 37°C: HSF in LB supplemented with Amp, Str, and NAL; AMD290 in LB supplemented with carbenicillin and diaminopimelic acid (DAP). Equal proportions of both strains were washed twice separately with LB containing DAP (for AMD290) or LB alone (for HSF). Final pellets were resuspended in 1 mL of LB containing DAP and mixed for conjugation. The mixture was conjugated for 5 hours at room temperature (25°C) on LB agar plates containing DAP.

After conjugation, cells were resuspended in LB and plated on LB agar with 100 μg/mL kanamycin to select for transposon mutants. After overnight incubation at 37°C, kanamycin-resistant (Kan^R) colonies were scraped together into 10 mL LB, and OD□□□ was measured. The mutant library was diluted to a starting OD□□□ of 0.2 in 60 mL LB with 100 μg/mL kanamycin and grown at 37°C to a final OD□□□ of 1.0. Glycerol was added to a final concentration of 15%, and 1 mL aliquots were stored at -80°C. Cells were collected for genomic DNA extraction.

To link random DNA barcodes to transposon insertion sites, genomic DNA was extracted from mutant library cell pellets using the DNeasy kit (Qiagen). Sequencing libraries compatible with Illumina were prepared according to established protocols^36,44^. Paired-end sequencing (150 bp) was performed with the NovaSeq system (Illumina). Mapping of transposon insertion sites and identification of corresponding DNA barcodes were carried out as previously described^36,44^. Due to the underlying construction parameters of the delivery system, this completed mutant library carries background Mu phage tracks. However, because the *E. coli* HSF host strain is naturally non-infective to classical dsDNA phages, we did not observe any confounding selective enrichment of conventional somatic dsDNA phage receptors during library processing, and all candidate hits in RB-TnSeq were subsequently confirmed.

### Competitive fitness experiments with RB-TnSeq library

For fitness assays, we followed the prior reported workflow^42,43^. Briefly, a 1 mL aliquot of the mutant library was thawed and inoculated into 25 mL of medium containing 100 μg/mL kanamycin, growing to OD□□□ of 0.6-0.8 at 37°C and 50rpm. Once the library reached mid-log phase, cell pellets were collected as a baseline reference for BarSeq (time-zero samples). The remaining cells were used for competitive fitness assays with different phages at multiplicities of infection (MOI) spanning 1 to 10.

For planktonic culture assays, the mutant library was diluted to an initial OD□□□ of 0.04 in 2X LB medium (350 μL), then mixed with an equal volume (350 μL) of phages diluted in SM buffer. Control assays without phages were prepared by replacing phages with SM buffer and 2X LB medium. Assays were performed in 48-well microplates (700 μL per well) and incubated in Tecan Infinite F200 readers with orbital shaking, taking OD□□□ readings every 15 min for 8 hours. Surviving cells were pelleted and stored at -80°C for subsequent genomic DNA extraction.

### BarSeq of RB-TnSeq fitness assays and data analysis

Genomic DNA from the RB-TnSeq library samples was extracted in a 96-well format with a QIAprep 96 Plus Kit (Qiagen). BarSeq PCR was performed using previously described protocols with minor modifications^42,43^. Briefly, BarSeq PCR was conducted in 50 μL volumes with 20 pmol of each primer and approximately 200 ng of template DNA. For NovaSeq runs, an equimolar mix of BarSeqV4_P7_S1 primers was used for demultiplexing alongside BarSeqV4_P5_S1 primers with additional sequences for sample verification. Equal volumes (5 μL) of each PCR reaction were pooled, purified, and eluted in 40 μL of water. BarSeq libraries were sequenced on an Illumina NovaSeq with 100 bp single-end reads,. RB-TnSeq fitness data were analyzed following previously established methods^36,44^.

### Deletion of F plasmid

The genes *repB* and *repE* of the F-plasmid were targeted to cure the F-plasmid from *E. coli* HSF. Oligos for crRNAs were designed as described previously and cloned into a plasmid expressing nuclease-active LbCas12a^85,98^. Sequence verified plasmids were transformed into *E. coli* HSF and selected on LB+Cam30 plates. A single colony was streaked onto an LB plate supplemented with Cam30 and 2 nM anhydrotetracycline (aTc), and incubated at 37°C overnight. Cleavage of the F-plasmid was driven by the nuclease-active LbCas12a; thus, ampicillin was omitted from the medium. A few colonies were patched into LB+Amp10 and LB plates to select for Amp^S phenotype, confirming curing of the F-plasmid. The selected Amp^S colony was grown overnight in LB and streaked to obtain single colonies. These colonies were patched onto LB+ Cam30 and LB plates to select for Cam^S colonies yielding an *E. coli* HS strain cured of the plasmid encoding the CRISPR-Cas system.

### Construction of mutant strains

Isogenic deletions of target genes (*dsbA*, *traA*) were generated using a recombineering technique based on the pSIM5 system. Primers were designed to amplify a kanamycin resistance (Kan^R) selection marker with 50 bp homology arms flanking target insertion sites, resulting in Kan^R insertions at each target gene^99^. PCR products were gel-purified using standard molecular biology methods and stored at -20°C until use. A temperature-sensitive pSIM5 recombineering vector was electroporated into *E. coli* HSF and cultured in a chloramphenicol-containing medium at 30°C. Cells were grown until the OD600 reached 0.1 at 30C at which point the culture was shifted to a shaker incubator maintained at 42°C to thermally induced lambda Red system. At an OD ∼0.6, cells were chilled on ice and made electrocompetent. PCR products with homology arms prepared as described above were electroporated, and after a 2-hour recovery growth step at 37°C, cells were plated on LB+Kan25 at 37°C. Kan^R colonies were selected and gene knockout were confirmed via Sanger sequencing of the resulting PCR product.

### Experimental validation of phage resistance phenotypes

To confirm phage resistance phenotypes observed in loss-of-function screens, efficiency of plating (EOP) assays were performed (Fig 4b, Supplementary Figs. S4-S6, S10-S11). All deletion strains and plasmids were verified through Sanger sequencing. For genetic complementation of *E. coli* HSF ΔdsbA, the plasmids pGRG25-Ptac::dsbA and pDM1 (kindly provided by Dr. Despoina Mavridou, University of Texas at Austin, USA) were used. The IPTG-inducible pGRG25-Ptac::dsbA system integrates dsbA into the host genome via Tn7 transposition^63^. To allow for IPTG induction, the lac repressor-encoded plasmid pDM1 was subsequently transformed into the integrated strain. Additionally, the native ampicillin resistance marker Amp^R of pGRG25 was replaced with a chloramphenicol resistance marker Cm^R to permit selection against the Amp^R-carrying F-plasmid.

Phage titers were determined by spot titration, where 2 μL of undiluted or 10-fold serially diluted phage solutions were spotted on solidified lawns of approximately 5 mL of 0.45% top agar inoculated with 100 μL of overnight bacterial culture and incubated overnight at 37°C. EOP values were calculated as the ratio of plaques formed on mutant or overexpression strains compared to parental strain HSF, based on at least three biological replicates.

### Evaluation of chemical inhibitors of DsbA

The capacity of chemical inhibitors to phenocopy genetic loss was evaluated by challenging the wild-type *E. coli* HSF with geraniol and 2-phenylthiophene (Sigma). Overnight cultures were cultivated in LB broth containing appropriate antibiotics at 37°C with shaking (200 rpm). Cultures were diluted 100-fold in media to reach OD□□□of approximately 0.05 (approximately 10□CFU/mL).

Stock solutions of each DsbA inhibitor were prepared at 1 M in DMSO and subsequently titrated in medium to establish a concentration gradient from 2 μM to 2,500 μM. DMSO and sterile LB were utilized as negative and positive controls, respectively, to account for baseline effects. Quantitative growth profiling determined that 63 μM represented the maximum sub-inhibitory concentration for both compounds, serving as the functional working dosage for phage replication assays. Bacterial growth was monitored in 96-well microtiter plates using a BioTek Logphase 600 reader to track absorbance at 600 nm following the addition of 10 CFU per well for 18 hours at 37°C.

### Comparative functional genomics

To identify conserved and strain-specific genes in the HSF strain and two *E. coli* lab strains, a comparison of chromosomally-encoded genes of *E. coli* strains HS (NCBI accession number CP092639), BW25113 (accession number CP122319), and C-3000 (ATCC 15597, downloaded from the ATCC website) was performed using OrthoFinder version 2.5.5 ^100^. Homologous sequences of the HSF capsular locus were identified by the Web-based BLASTN search tool at NCBI. Prophages were predicted by geNomad^101^, and phage defense systems were analyzed by DefenseFinder v2^81^. Multiple alignment of *E. coli* genomes was constructed by progressiveMauve^102^.

Targeted searches for K47-capsule depolymerase domains were conducted within the genomes of *E. coli* dsDNA phages HSDP1 and HSDP2 by retrieving tail protein-coding genes and identifying homologous domains with PaperBLAST^103^ and the Conserved Domains Database v3.20 (CDD)^104^. The similarity of phage HSDP2 relative to other Tequatrovirus phages was estimated by calculating the pairwise proteomic equivalent quotient (PEQ)^105^. PEQ values calculated against classical phages T2 and T4 were 0.865 and 0.822, respectively. Since no annotated depolymerase domain was found in any of the HSDP2 tail fiber proteins, its gp37 distal long tail fiber protein (ZHKDYPRP_CDS_0080) was aligned with representative gp37 protein sequences from other T4-like phages to search for unique domains. Proteins encoded in the tail locus of phage HSDP1 (PCOOMCAM_CDS_0037 to PCOOMCAM_CDS_0051) were systematically searched with PaperBLAST to identify putative depolymerase domains that would explain its ability to bypass HSF’s capsule. Protein sequence alignment was visualized with the ggmsa v1.0.3 R package^106^. The 3D structures of the tail proteins were modeled with AlphaFold3^107^ and annotated in ChimeraX v1.10^108^.

### Extraction and purification of the surface glycolipids

The analytical and compositional analysis was performed with minor modifications from prior work^109–111^. The bacterial pellets of wild-type (WT HSF) (2.8 g) and mutant (HSF Δ01649) (2.3 g) were uniformly suspended in 15 mL of dH2O, prewarmed to 68 °C, and extracted with 15 mL of prewarmed liquefied phenol at 68 °C for 20 min following the Westphal method^112^. The extracts were cooled on ice and centrifuged at 5000 x g for 20 min at 4 °C. The upper water layer was carefully transferred, and an equal amount of deionized water was added to each bacterial pellet, and the extraction was repeated two more times. The combined water phases and the phenol layer were dialyzed against three changes of deionized water for three consecutive days using 12-14 kDa MWCO dialysis tubing at 4°C. The dialysates were collected and freeze-dried.

Because phenol-phase recovery was minimal (<2 mg), processing was continued only with the water layer extracts. The crude extracts were washed three times with chilled 95% (vol./vol.) ethanol, followed by brief sonication and centrifugation for 20 min at 3000 x g. The nucleic acids and proteins were digested overnight with 50.4 U/mL Benzonase (Millipore Sigma/Merck KGaA) at 37 °C, followed by overnight digestion with 0.82 U/mL Proteinase K (VWR) at 37 °C. Both reactions were facilitated by gentle agitation at 100 rpm. The degraded nucleotides, peptides, and buffer were dialyzed out against three changes of dH2O for three consecutive days using 12-14 kDa MWCO dialysis tubing at 4 °C, and the dialysates were freeze-dried. Finally, the glycolipids were centrifuged at approx. 100,000 x g for 16 h at 4 °C using an Ultra OptimaTM L-90K ultracentrifuge. Ultimately, 6.0 mg of ultracentrifugation supernatant and 2.0 mg of pellets were recovered from *E. coli* HSF WT, and 1.0 mg of ultracentrifugation supernatant and 1.9 mg of pellets were obtained from *E. coli* HSF Δ01649 mutant, respectively.

### DOC-PAGE

The fractions recovered from ultracentrifugation (1 µg) were resolved by PAGE (4% stacking gel and 18% resolving gel) in the presence of the deoxycholic acid buffer^111^. Resulting bands were visualized with a silver stain reagent kit (Bio-Rad).

### Composition analysis

The glycosyl and fatty acid composition was determined by conversion to per-*O*-trimethylsilyl (TMS) methyl glycosides and fatty acid methyl ester (FAME) and TMS-FAME derivatives for hydroxylated fatty acids, as described earlier^109,110^. Briefly, 200 µg of the sample was mixed with 10 µg of the inositol internal standard and freeze-dried. Samples were converted to methyl ethers of monosaccharides and fatty acid methyl esters by methanolysis in 1 M HCl-methanol at 80°C for 18 h, then dried under a stream of dry air. Re-*N*-acetylation of amino sugars was carried out using 200 µl of methanol, 100 µl of pyridine, and 100 µl of acetic anhydride at 80 °C for 1 h. After evaporation of the reagents, the samples were *O*-trimethylsilylated using 300 µl of Tri-Sil HTP reagent (Thermo) at 80□for 20 min, followed by gentle evaporation. The derivatized samples were dissolved in hexane and analyzed by GC-MS (Agilent 7890A GC interfaced to a 5975C MSD), using a Supelco Equity-1 fused silica capillary column (30 m × 0.25 mm ID). After 2 min at the initial temperature of 80 °C, the temperature was increased at 2 °C/min to 200 °C with a 2 min hold, then at 30 °C/min to 250 °C with a 5 min hold.

### cryo-ET data collection and processing

Overnight cultures of *E. coli* strains (WT HSF strain or Δ01649_HSF) were inoculated at a 1:100 ratio into 30 ml LB and grown until an OD550 of 0.45 was reached. Then, 50 µl of *E. coli* culture was mixed with 20 µl MS2 (10¹□PFU/ml) and incubated at 37 °C for 20 min before addition of 10 nm fiducial gold (EMS).

A 4 µl aliquot of the sample was applied to Quantifoil R3.5/1 grids (copper, 300 mesh) and vitrified using a Leica GP2. Data collection was performed using a Titan Krios G4 (ThermoFisher) equipped with a Gatan K3 camera at a nominal magnification of 19,500×, yielding a pixel size of 4.5 Å/pixel. Each tilt image was acquired with a total exposure of 2.5 s over a range of ±54°, using a 3° tilt increment, yielding a cumulative dose of ∼120□e^−^/Å.

The tilt series were aligned using MotionCor2^113^ and reconstructed with IMOD^114^ to produce tomograms binned by four. Tomograms were denoised using IsoNet^115^. MS2 particles were annotated using PyTom template matching^116^ with EMD-3403^117^. F pili were manually annotated in IMOD(^114^), and putative T4SS machinery was manually annotated using ArtiaX^118^ and visualized in ChimeraX^119^.

## Data Availability

Assembled genome sequences for all phages characterized in this study have been deposited in NCBI GenBank under accession numbers (submitted).

## Acknowledgements

This material by the Biopreparedness Research Virtual Environment (BRaVE) Phage Foundry at Lawrence Berkeley National Laboratory is based upon work supported by the U.S. Department of Energy, Office of Science, Office of Biological & Environmental Research under contract number DE-AC02-05CH11231. The work conducted by the U.S. Department of Energy Joint Genome Institute (https://ror.org/04xm1d337), a DOE Office of Science User Facility, is supported by the Office of Science of the U.S. Department of Energy operated under Contract No. DE-AC02-05CH11231. The work at the Complex Carbohydrate Research Center was supported in part by the U.S. Department of Energy, Office of Science, Basic Energy Sciences, Chemical Sciences, Geosciences and Biosciences Division, under award DE-SC0015662 to the DOE Center for Plant and Microbial Complex Carbohydrates at the CCRC and NIH R24GM137782 award to the National Glycoscience Resource-CCRC Service and Training to P.A. This work at Texas A&M University was supported by the Center for Phage Technology, the NIH R01GM141659, the TAMU ADM grant. For cryo-ET data collection, the authors acknowledge the Texas A&M University Laboratory for Biomolecular Structure and Dynamics, jointly supported by the Department of Biochemistry and Biophysics, AgriLife, Texas A&M University, and the Cancer Prevention and Research Institute of Texas (CPRIT). We are grateful to Tagide deCarvalho at the Keith R. Porter Imaging Facility (University of Maryland, Baltimore County) for assistance with transmission electron microscopy imaging. Molecular graphics and analyses performed with UCSF ChimeraX, developed by the Resource for Biocomputing, Visualization, and Informatics at the University of California, San Francisco, with support from National Institutes of Health R01-GM129325 and the Office of Cyber Infrastructure and Computational Biology, National Institute of Allergy and Infectious Diseases. This work was also supported by the Laboratory Directed Research and Development (LDRD) program at Lawrence Berkeley National Laboratory (to S.R and V. K. M).

## Supplementary Figures

**Supplementary Figure S1.** Genogroup stratification of *Leviviricetes* phage isolates via diagnostic RT-PCR multiplexing

**Supplementary Figure S2.** Comparative genomic maps and structural annotations of isolated ssRNA phages.

**Supplementary Figure S3.** Transmission electron microscopy (TEM) morphological analysis of isolated single-stranded RNA phage virions

**Supplementary Figure S4.** Titration of full panel of phages on *E. coli* HSF.

**Supplementary Figure S5.** Validation spot testing and efficiency of plating (EOP) assays.

**Supplementary Figure S6.** Effect of *traD* deletion across ssRNA genogroups

**Supplementary Figure S7.** Chemical inhibition of host DsbA in *E. coli* HSF by geraniol phenocopies genetic loss during ssRNA phage infection

**Supplementary Figure S8.** Chemical inhibition of DsbA substrate-binding groove in *E. coli* HSF by 2-phenylthiophene (2-PTP) phenocopies genetic loss during ssRNA phage infection

**Supplementary Figure S9**. Genomic fragment and gene-level fitness scores from Dub-seq expression screening under R17 phage selection

**Supplementary Figure S10.** Host capsule expression in *E. coli* HSF restricts infection across a broad collection of 156 dsDNA phages.

**Supplementary Figure S11.** Capsule disruption does not impact ssRNA phage infection.

**Supplementary Figure S12.** Chemical characterization of cell surface glycolipids isolated from wild-type *Escherichia coli* HSF (a) and the capsule-deficient Δ*01649* mutant (b).

## Supplementary Tables

**Supplementary Table S1**. Bacteriophages utilized in this study, their isolation sources, and propagation parameters.

**Supplementary Table S2**. Bacterial strains, mutants, plasmids, and oligonucleotide primers used in this work.

**Supplementary Table S3**. Quantitative host gene fitness scores from RB-TnSeq screens under ssRNA phage selection.

**Supplementary Table S4**. Quantitative host fitness scores from Dub-seq screens under ssRNA phage selection.

**Supplementary Table S5.** Quantitative host gene fitness scores from RB-TnSeq screens under filamentous ssDNA phage selection.

**Supplementary Table S6**. Comparative genomic and bioinformatic analysis of the *Escherichia coli* HSF capsular polysaccharide locus.

**Supplementary Table S7.** Quantitative host gene fitness scores from RB-TnSeq screens under dsDNA phage selection.

**Supplementary Table S8.** Glycosyl residues detected in the glycolipid fraction recovered from ultracentrifugation sediments of *E. coli* HSF WT and Δ01649 mutant.

**Supplementary Table S9.** Fatty acid residues detected in the glycolipid fraction recovered from ultracentrifugation sediments of *E. coli* HSF WT and Δ01649 mutant.

