## Supplementary material for "Genome-wide characterization of host factors involved in single-stranded RNA and DNA phage infection pathways": Supplmentary Figures

**Supplementary Figures**

**Supplementary Figure S1.** Genogroup stratification of *Leviviricetes* phage isolates via diagnostic RT-PCR multiplexing

**Supplementary Figure S2.** Comparative genomic maps and structural annotations of isolated ssRNA phages.

**Supplementary Figure S3.** Transmission electron microscopy (TEM) morphological analysis of isolated single-stranded RNA phage virions

**Supplementary Figure S4.** Titration of full panel of phages on *E. coli* HSF.

**Supplementary Figure S5.** Validation spot testing and efficiency of plating (EOP) assays.

**Supplementary Figure S6.** Effect of *traD* deletion across ssRNA genogroups

**Supplementary Figure S7.** Chemical inhibition of host DsbA in *E. coli* HSF by geraniol phenocopies genetic loss during ssRNA phage infection

**Supplementary Figure S8.** Chemical inhibition of DsbA substrate-binding groove in *E. coli* HSF by 2-phenylthiophene (2-PTP) phenocopies genetic loss during ssRNA phage infection

**Supplementary Figure S9**. Genomic fragment and gene-level fitness scores from Dub-seq expression screening under R17 phage selection

**Supplementary Figure S10.** Host capsule expression in *E. coli* HSF restricts infection across a broad collection of 156 dsDNA phages.

**Supplementary Figure S11.** Capsule disruption does not impact ssRNA phage infection.

**Supplementary Figure S12.** Chemical characterization of cell surface glycolipids isolated from wild-type *Escherichia coli* HSF (a) and the capsule-deficient *Δ01649* mutant (b).

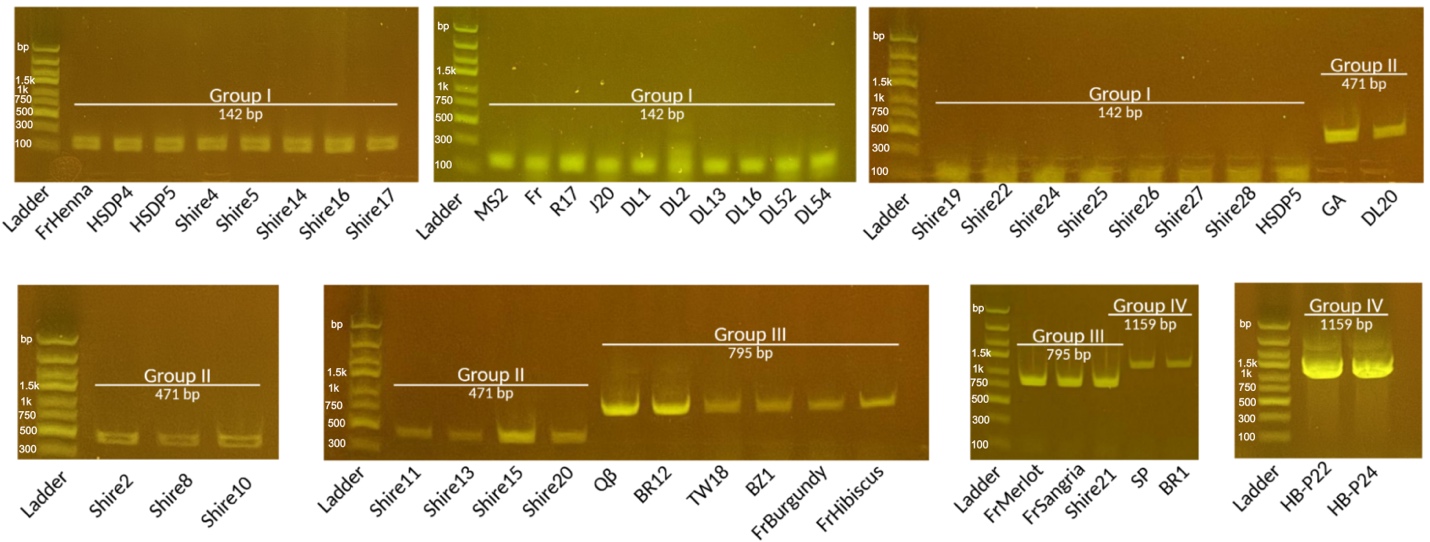

**Supplementary Figure S1. Genogroup stratification of *Leviviricetes* phage isolates via diagnostic RT-PCR multiplexing**. Gel-electrophoretic profile showing the classification of the 46 single-stranded RNA (ssRNA) phages characterized in this work into their respective genogroups. Following reverse transcription, genogroup-specific primer sets selectively amplify target regions within the viral genomes, yielding distinct, diagnostic amplicon sizes. Resulting DNA amplicons for each of the 46 isolates were resolved on an E-Gel system alongside an E-Gel DNA 1Kb Express Ladder (Thermo Fisher Scientific, catalogue number 10488090) for molecular weight verification. Extended assay details and primer configurations are described in the Methods.

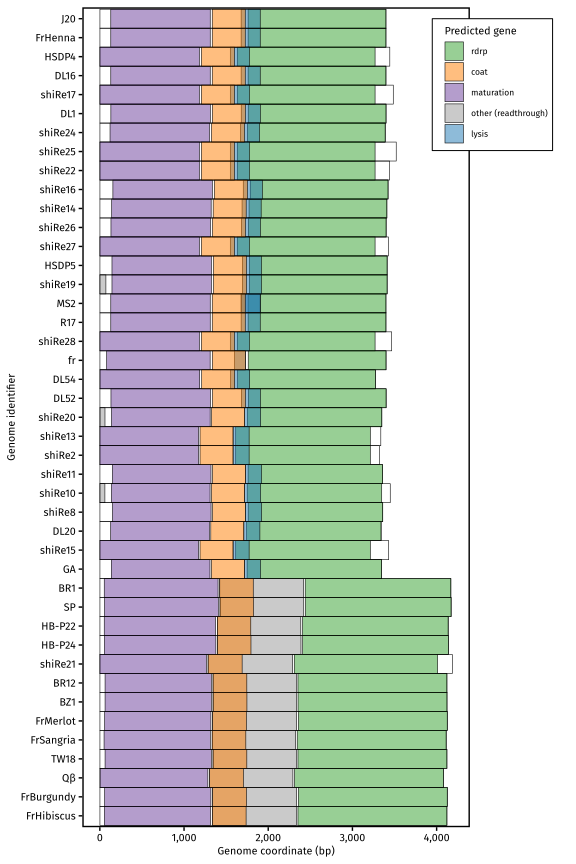

**Supplementary Figure S2. Comparative genomic maps and structural annotations of isolated ssRNA phages**. To-scale schematic representation of the ssRNA phage genome architectures. Open reading frames (ORFs) are depicted as directional horizontal blocks and color-coded based on their functional annotations, including maturation (mat), capsid coat (coat), lysis (sgl), and RNA replicase (rdrp, rep) genes.

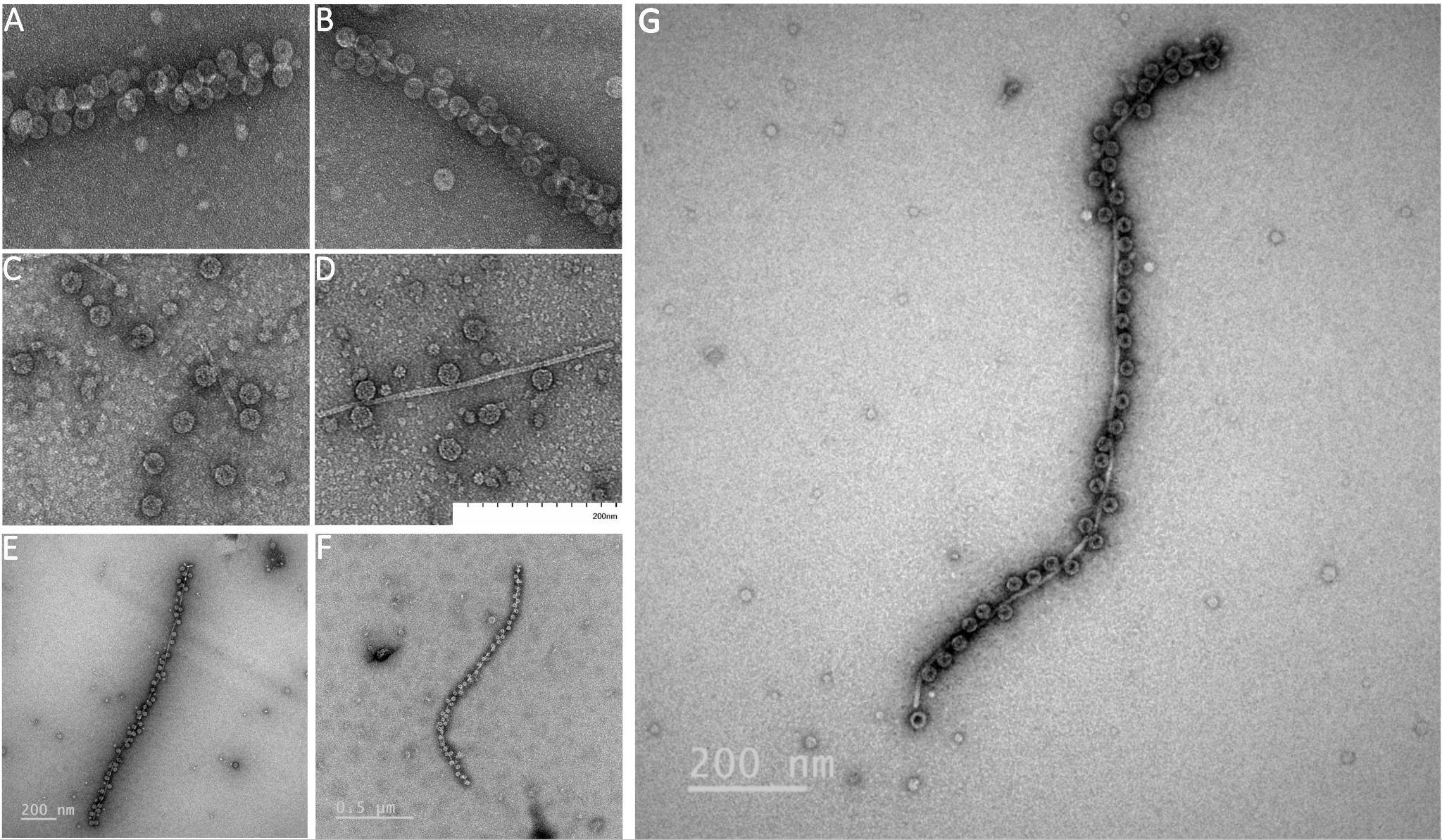

**Supplementary Figure S3. Transmission electron microscopy (TEM) morphological analysis of isolated single-stranded RNA phage virions**. Representative negative-stained transmission electron micrographs of selected ssRNA phages showcasing the characteristic small, icosahedral capsid associated with F pili. Panels display: (A) MS2 (B) GA (C) Qβ (D) SP (E) Shire17 (capsid diameter 27.04 ± 0.77 nm, pilus diameter 9.13 ± 0.73 nm) (F) Shire20 (capsid diameter 27.33 ± 1.2 nm, pilus 8.26 ± 1.00 nm) (G) BR1(capsid diameter 25.5 ± 0.44 nm, pilus diameter 9.32 ± 1.68 nm). Values are reported as mean ± SD. All virions display the characteristic small, non-enveloped, isometric morphology typical of members of the class Leviviricetes. Details regarding sample negative staining protocols and magnification levels are described in the Methods.

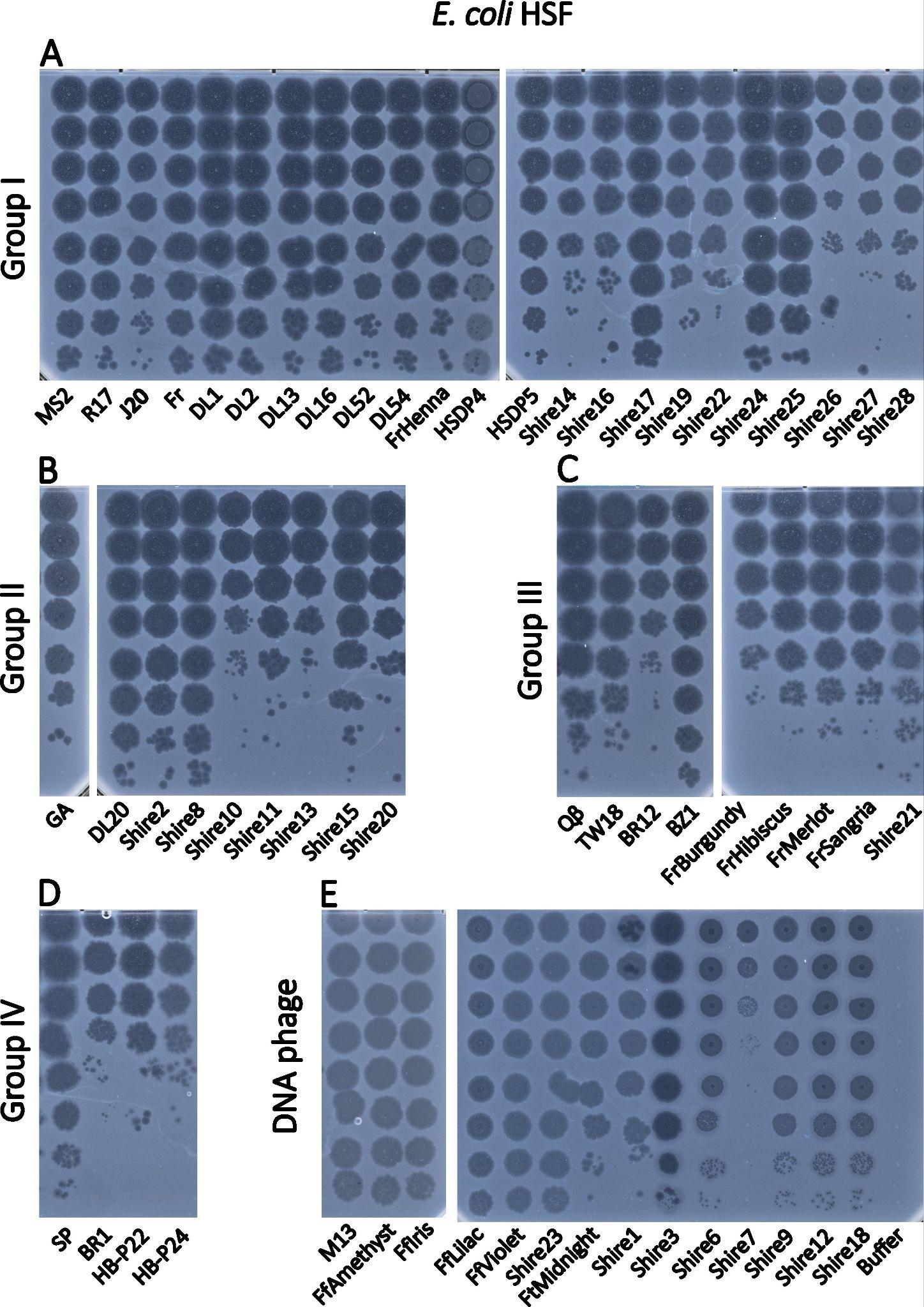

**Supplementary Figure S4. Titration of full panel of phages on *E. coli* HSF.** Viral titrations of the ssRNA groups I (A), II (B), III (C), IV (D), and (E) ssDNA and dsDNA phage collections on wild-type *E. coli* HSF (Supplementary Table S1).

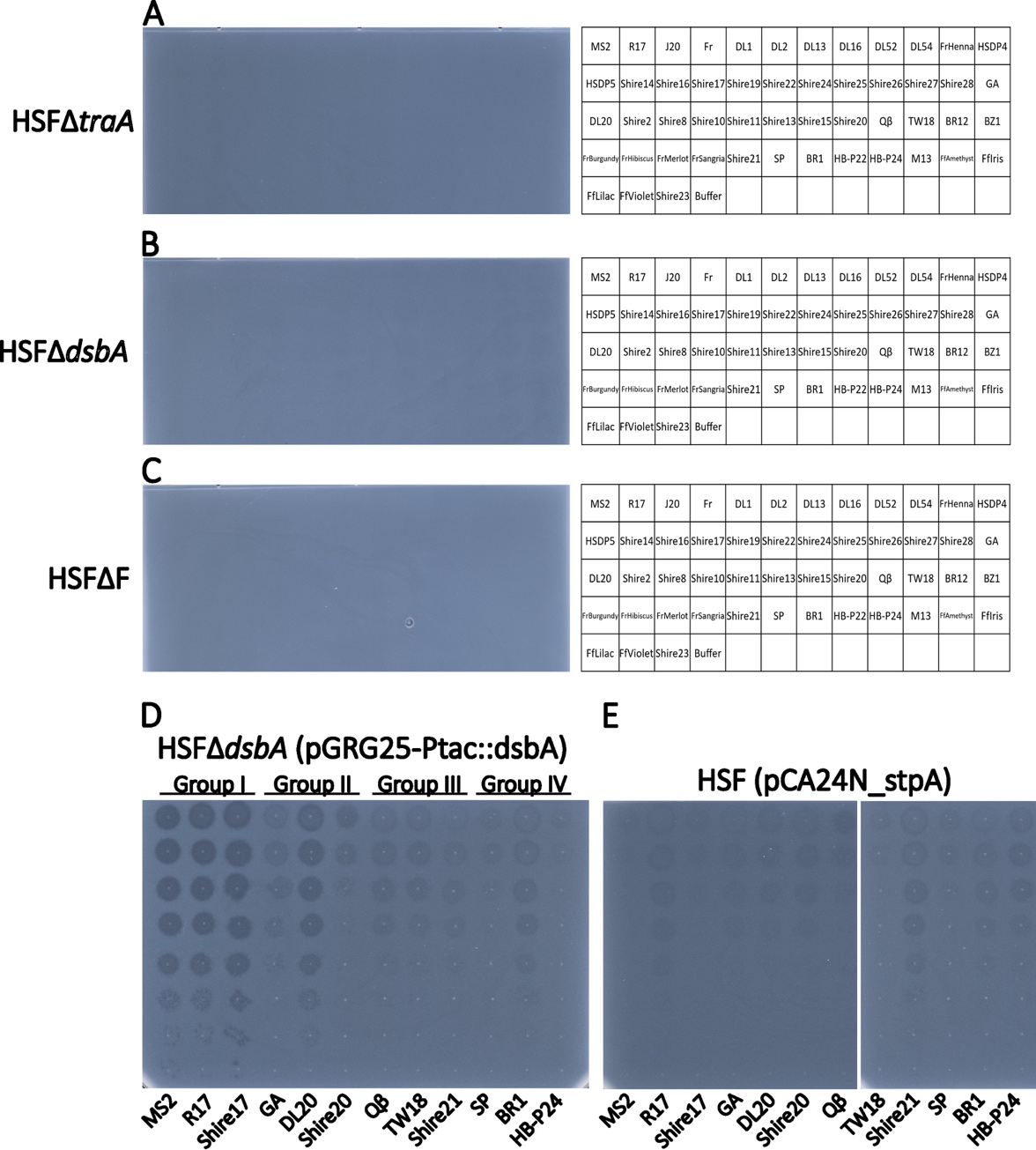

**Supplementary Figure S5. Validation spot testing and efficiency of plating (EOP) assays.** Targeted host susceptibility screenings across clean genetic deletion strains to confirm essential entry determinants. Direct spot assays against HSF strain (A) Δ*traA* (pilin subunit mutant), (B) Δ*dsbA* (periplasmic disulfide bond formation factor), and (C) the plasmid-cured F⁻ derivative demonstrate a universal block to infection across the entire single-stranded viral panel when these machinery elements are disrupted. (D) A degree of susceptibility is restored to the HSFΔ *dsbA* strain when complemented. (E) Infection is inhibited across the panel when plated on the HSF host overexpressing RNA chaperone *stpA*. Titration of full panel of ssRNA phages on WT *E. coli* HSF is given on Supplementary Figure S4.

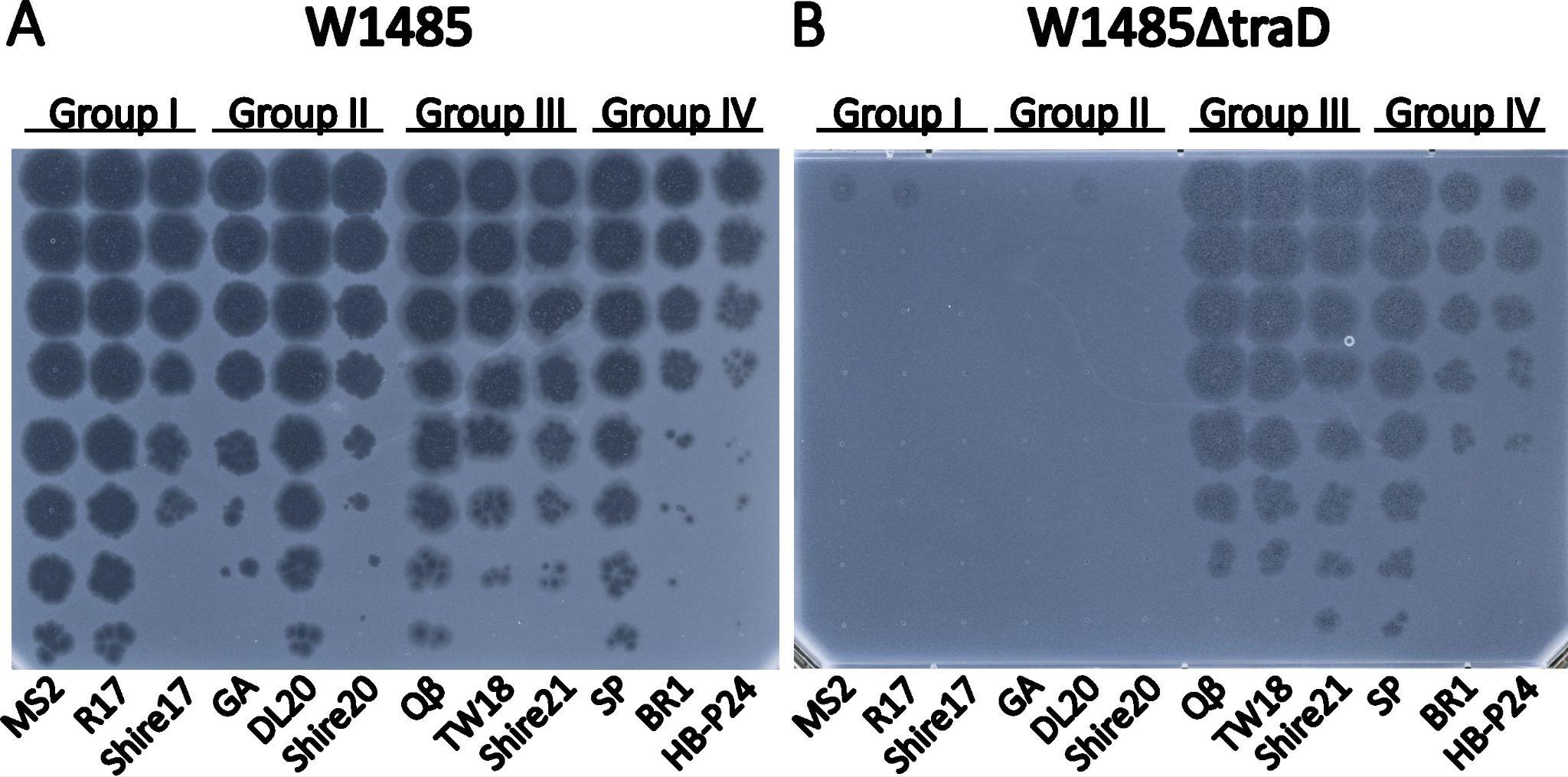

**Supplementary Figure S6. Effect of *traD* deletion across ssRNA genogroups.** Quantitative phage titrations demonstrating the genogroup-specific requirements of the F-pilus coupling factor TraD. Three representative phage isolates selected from each distinct genogroup were evaluated on (A) the F⁺ reference strain *E. coli* W1485 and (B) its corresponding conjugate mutant strain *E. coli* W1485 Δ*traD*.

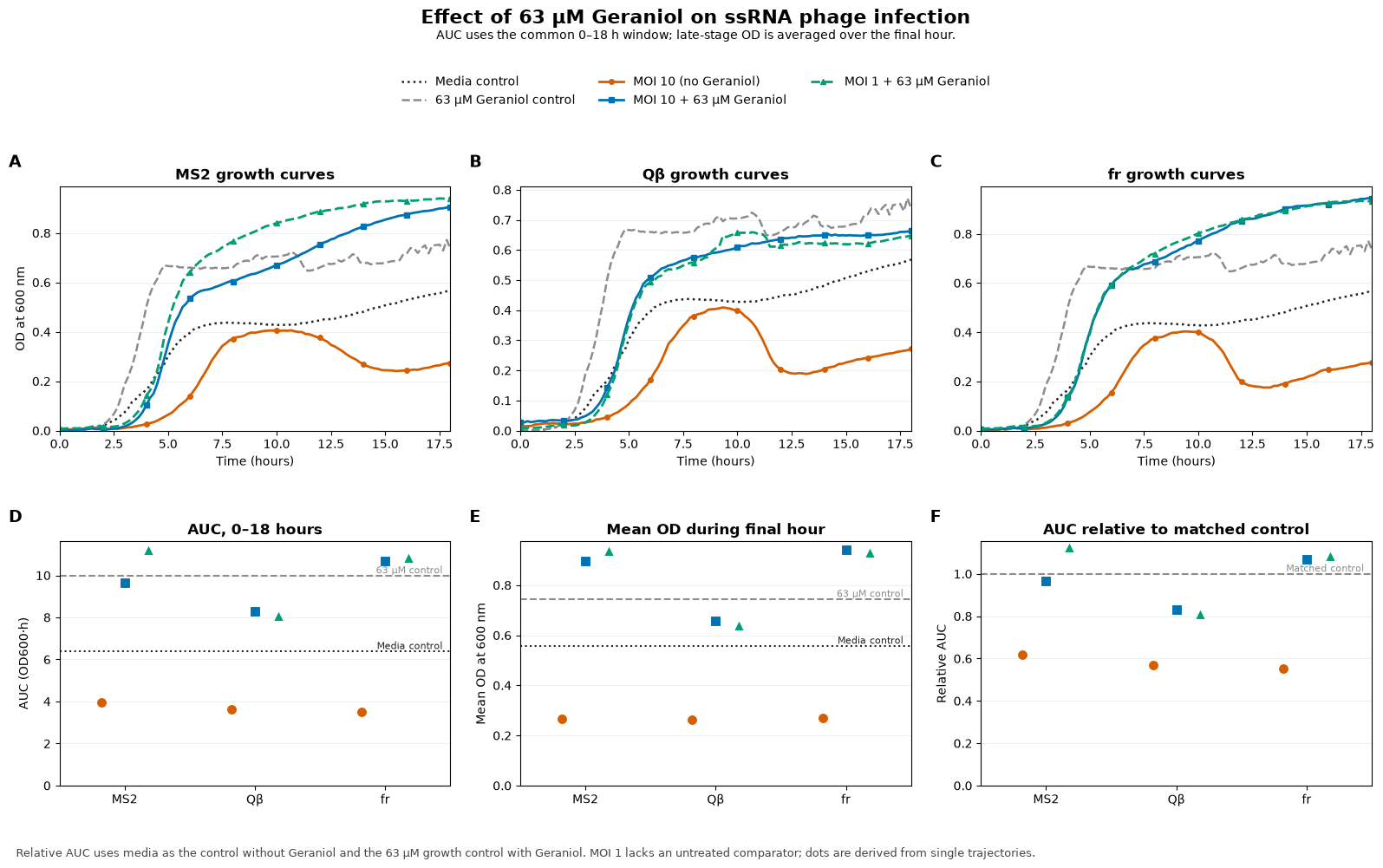

**Supplementary Figure S7. Chemical inhibition of host DsbA in *E. coli* HSF by geraniol phenocopies genetic loss during ssRNA phage infection** (A–C) Bacterial growth, measured as OD₆₀₀, was monitored for 18 h in the presence of MS2 (A), Qβ (B), or fr (C). Conditions were: phage at MOI 10 without geraniol (orange circles), phage at MOI 10 with 63 µM geraniol (blue squares), and phage at MOI 1 with 63 µM geraniol (green triangles). The bacterial media control without phage or geraniol (black dotted line) and the 63 µM geraniol growth control without phage (gray dashed line) are shown for reference. (D) AUC calculated by trapezoidal integration over the common 0–18 h interval. (E) Mean OD₆₀₀ during the final hour of the experiment. Horizontal dotted and dashed lines in D and E denote the media and 63 µM geraniol controls, respectively. (F) AUC relative to the matched no-phage growth control. Untreated MOI 10 conditions were normalized to the media control, whereas geraniol-treated conditions were normalized to the 63 µM geraniol control; a relative AUC of 1 denotes growth equivalent to the appropriate no-phage control. Relative AUC values for untreated MOI 10 cultures ranged from 0.55 to 0.62. With geraniol, relative AUC values were 0.97 and 1.12 for MS2, 0.83 and 0.81 for Qβ, and 1.07 and 1.09 for fr at MOI 10 and MOI 1, respectively. Thus, geraniol-treated cultures approached or exceeded the matched growth control for MS2 and fr, while the recovery of growth was slightly lower for Qβ.

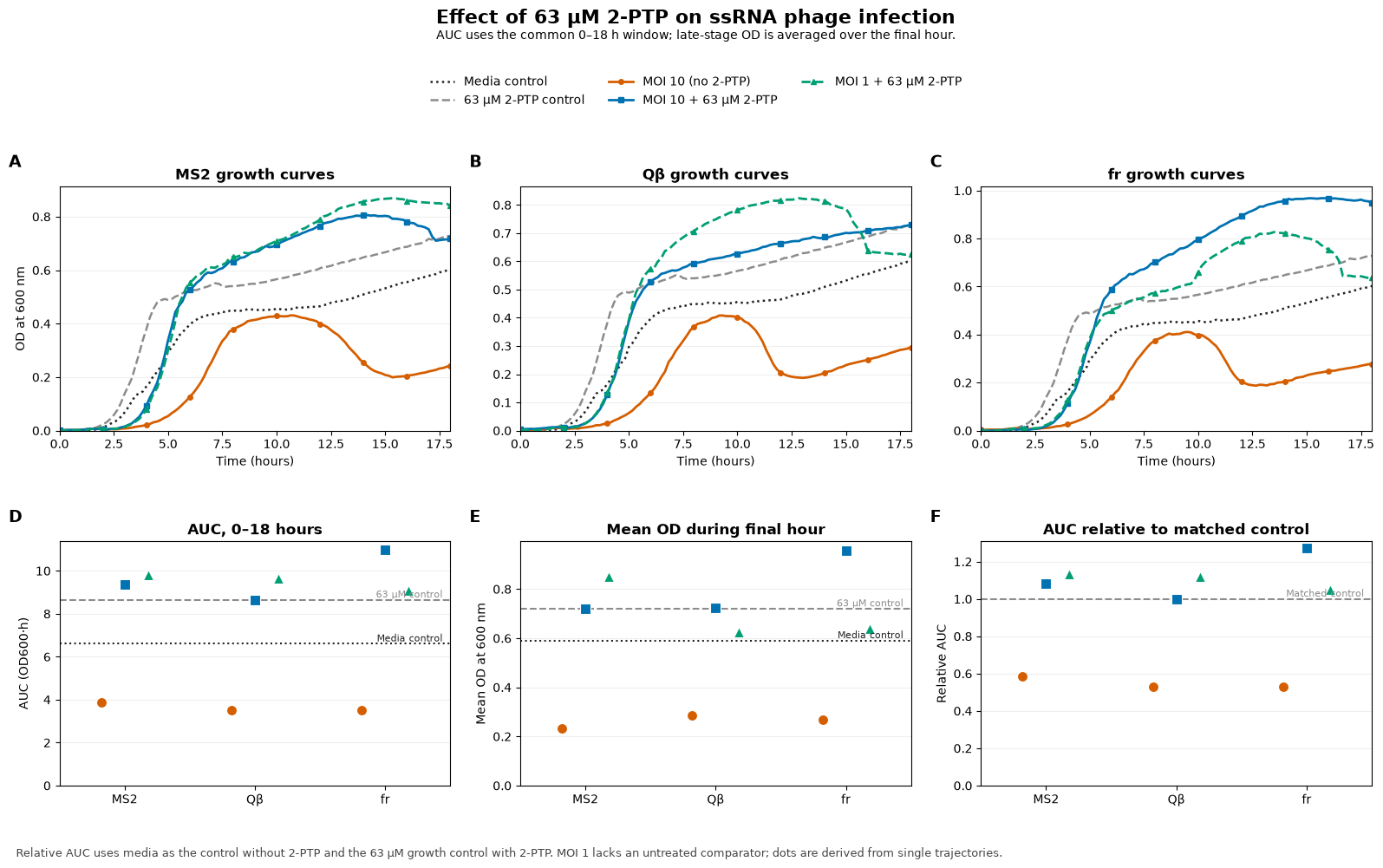

**Supplementary Figure S8. Chemical inhibition of DsbA substrate-binding groove in *E. coli* HSF by 2-phenylthiophene (2-PTP) phenocopies genetic loss during ssRNA phage infection** (A–C) Bacterial growth, measured as optical density at 600 nm (OD₆₀₀), was monitored for 18 h in the presence of the ssRNA phages MS2 (A), Qβ (B), or fr (C). Conditions were phage at a multiplicity of infection (MOI) of 10 without 2-PTP (orange circles), phage at MOI 10 with 63 µM 2-PTP (blue squares), and phage at MOI 1 with 63 µM 2-PTP (green triangles). The bacterial media control without phage or 2-PTP (black dotted line) and the 63 µM 2-PTP growth control without phage (gray dashed line) are shown for reference. (D) Area under the OD₆₀₀ growth curve (AUC), calculated by trapezoidal integration over the common 0–18 h interval. (E) Mean OD₆₀₀ during the final hour of the experiment. In D and E, horizontal dotted and dashed lines indicate the media and 63 µM 2-PTP controls, respectively. (F) AUC normalized to the matched no-phage growth control. Untreated MOI 10 conditions were normalized to the media control, whereas 2-PTP-treated conditions were normalized to the 63 µM 2-PTP control; a value of 1 therefore indicates growth equivalent to the corresponding no-phage control. Untreated MOI 10 cultures had relative AUC values of 0.53–0.59, consistent with phage-associated growth suppression. In the presence of 2-PTP, relative AUC values increased to approximately 1.00–1.27 at MOI 10 and 1.05–1.14 at MOI 1 across the three phages, indicating preservation of bacterial growth under these assay conditions.

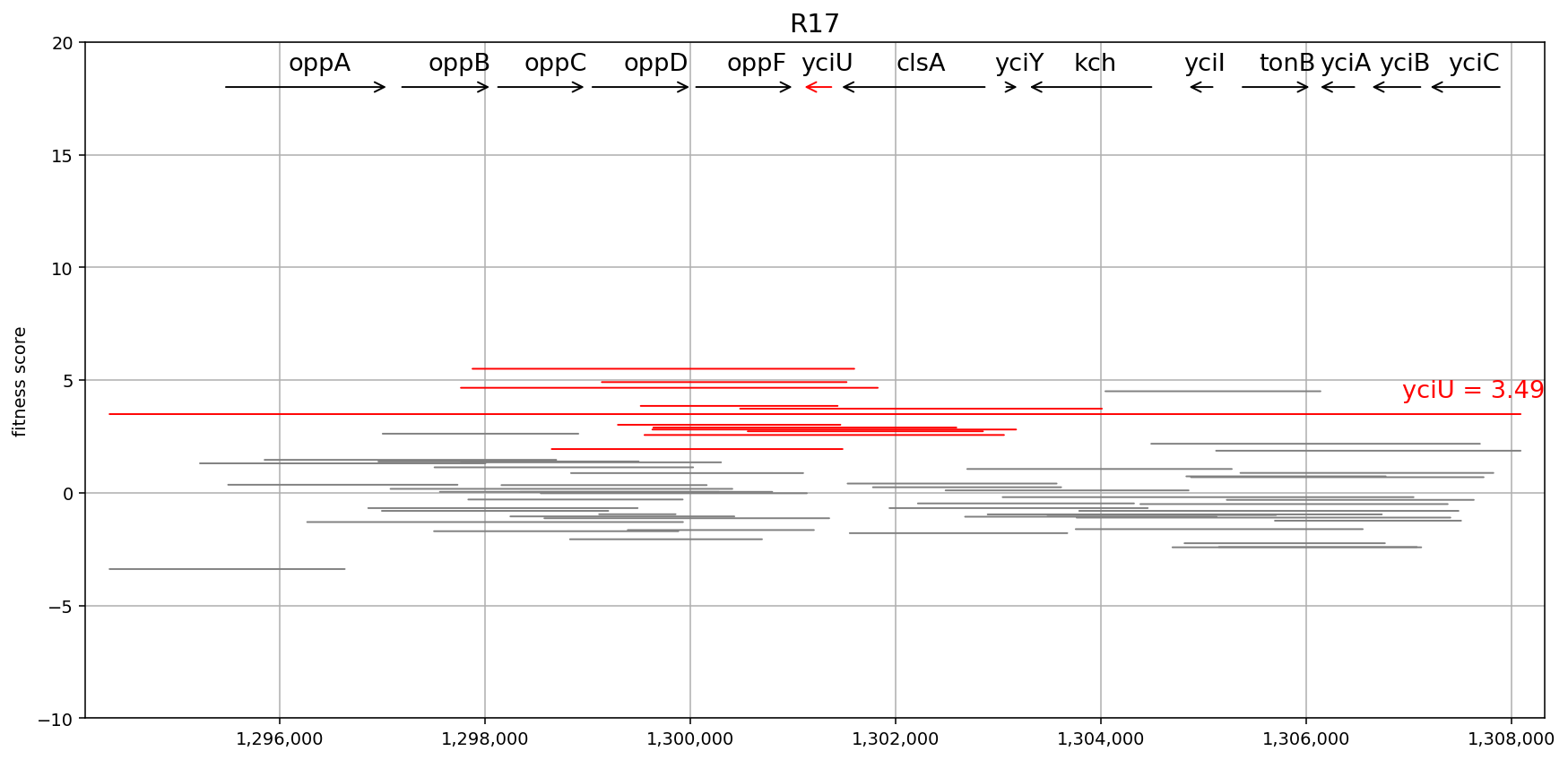

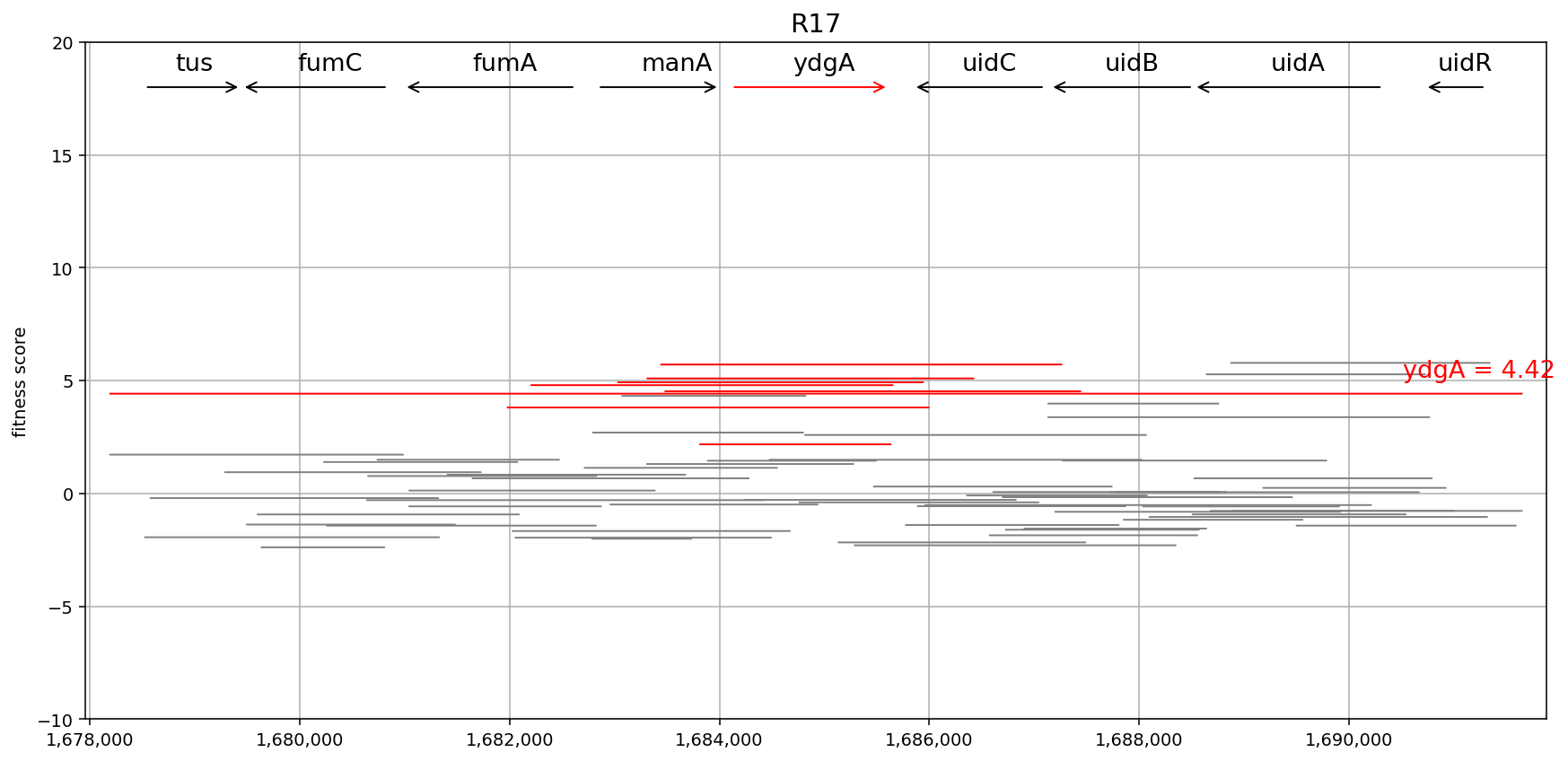

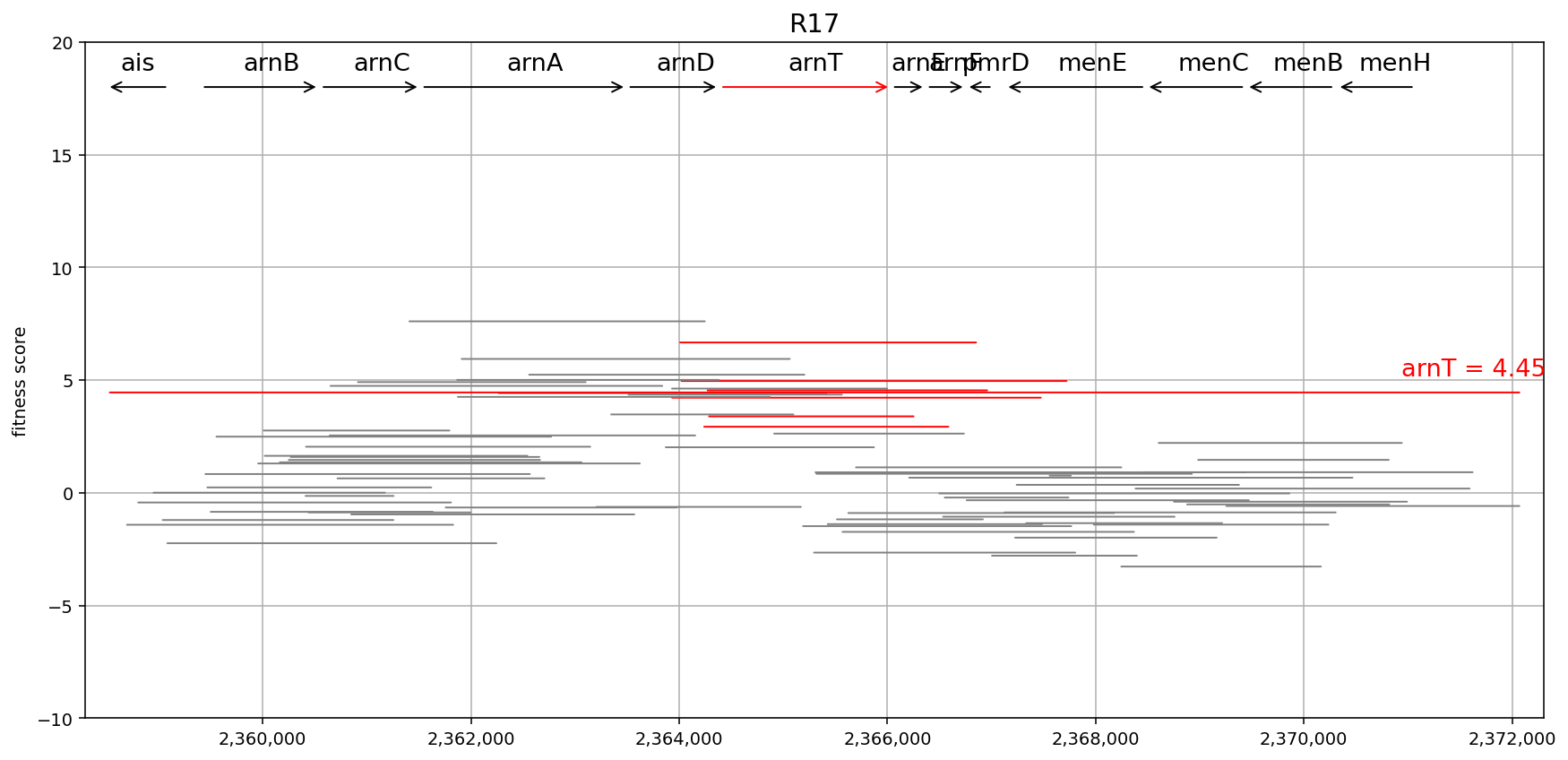

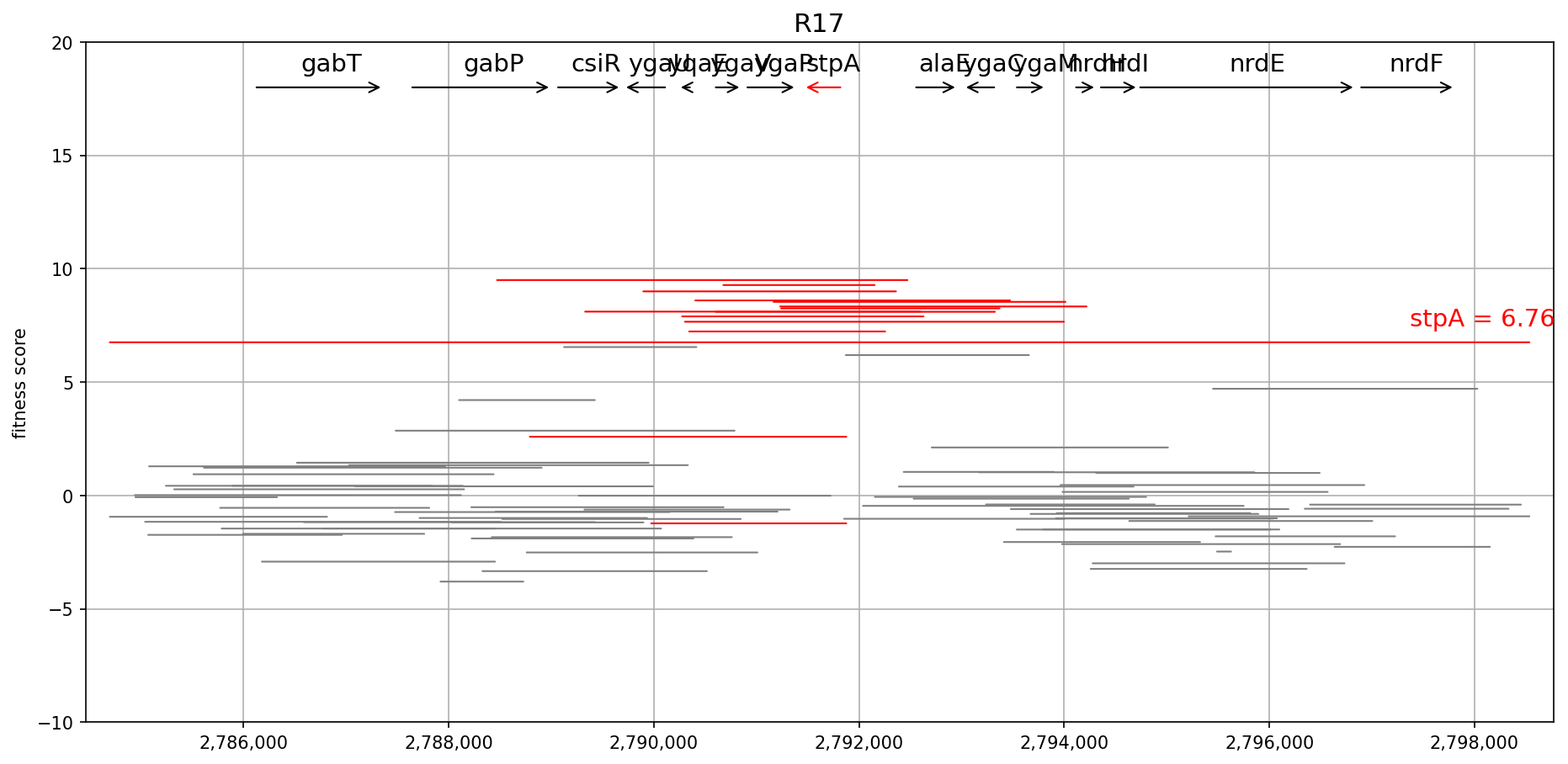

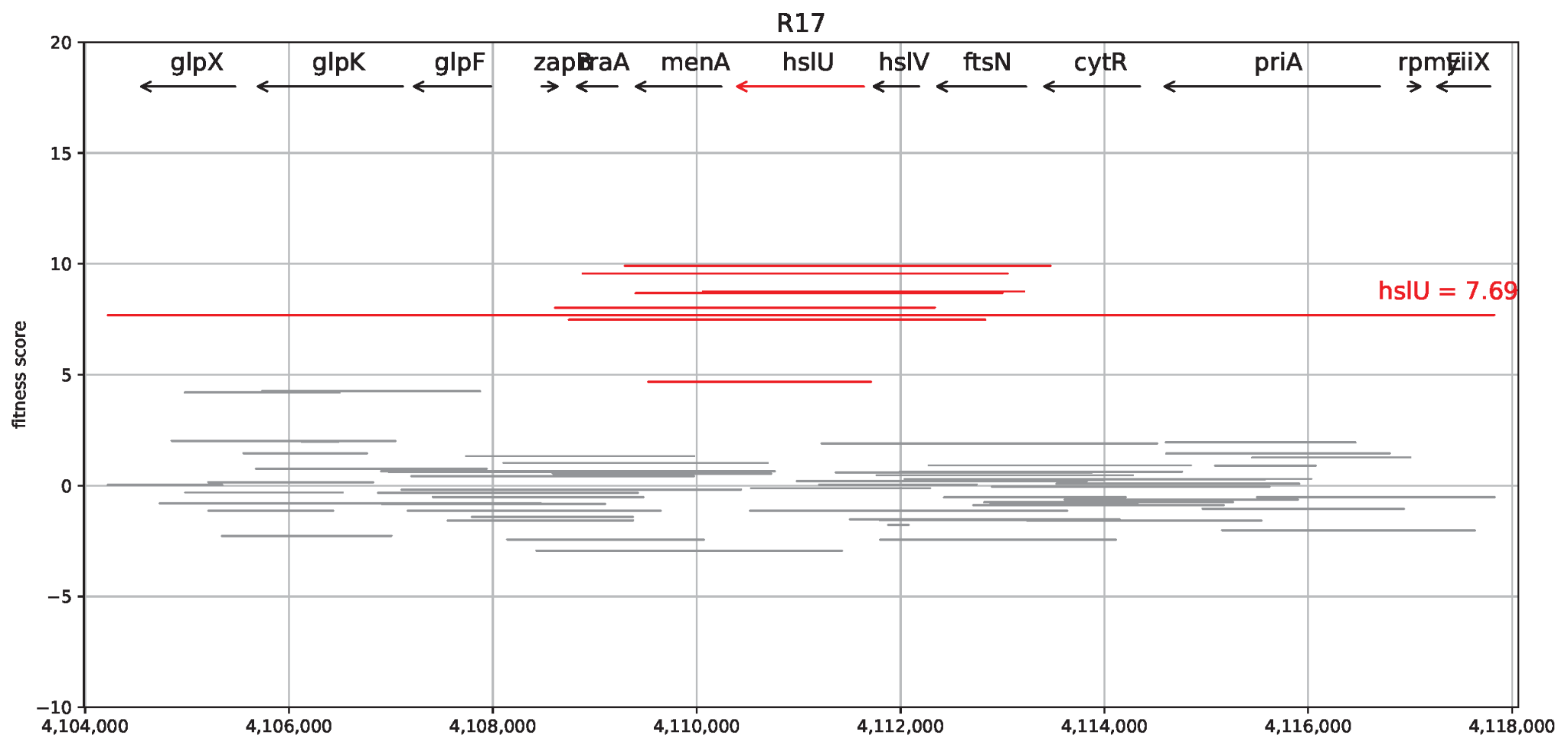

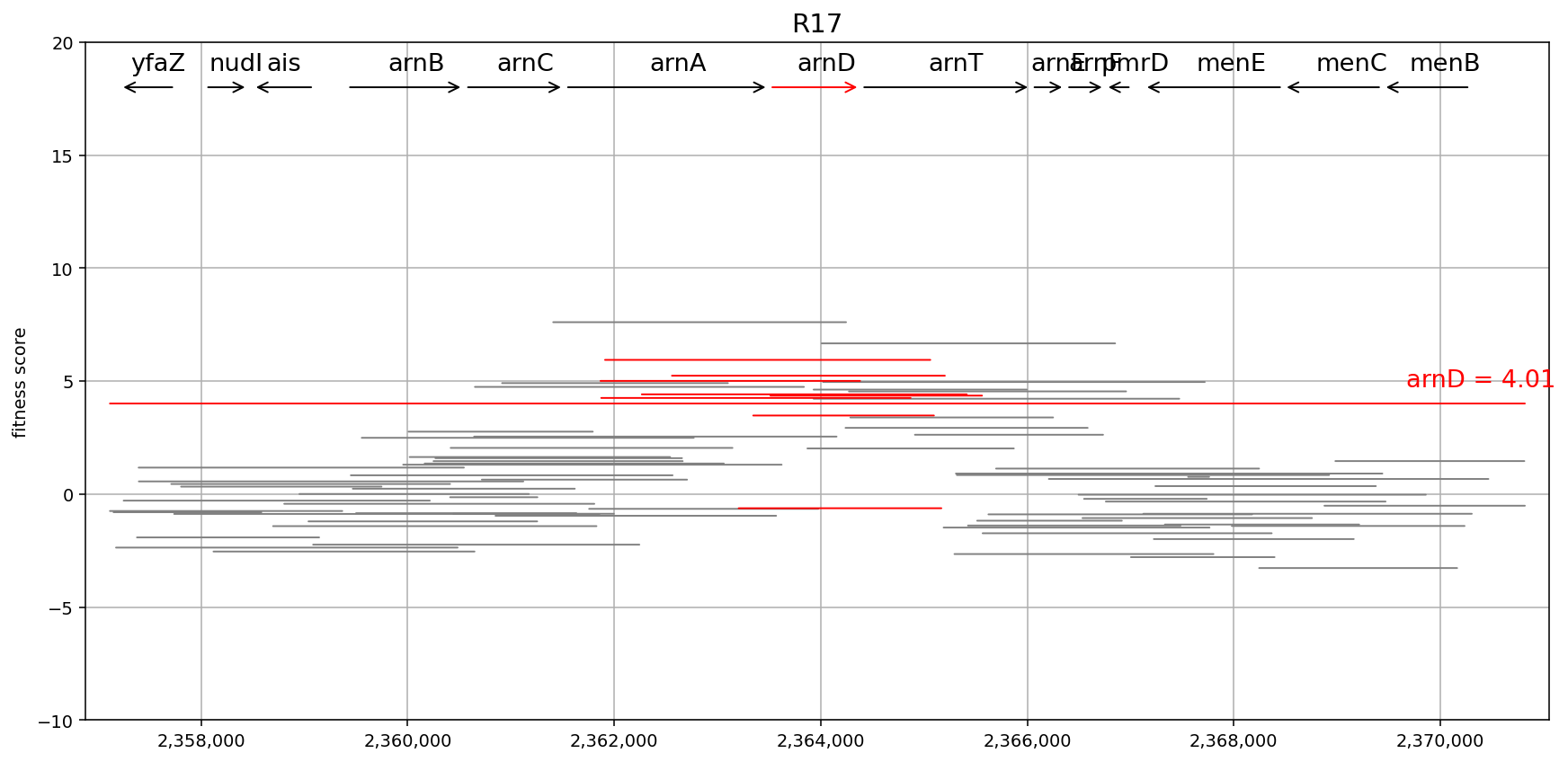

**Supplementary Figure S9. Genomic fragment and gene-level fitness scores from Dub-seq expression screening under R17 phage selection**. Locally zoomed-in genomic coordinate maps tracking individual Dub-seq fragment trajectories across locus regions surrounding high-confidence hits: *hslU*, *stpA*, *arnT*, *arnD*, *yciU*, and *ydgA*. The vertical axis maps the calculated strain fitness score under selective pressure from the ssRNA R17 phage. Each horizontal line corresponds to a distinct cloned genomic library fragment. Fragments that entirely span the full-length open reading frame of the top-scoring target gene are highlighted in red to validate gene-context specific phenotypes.

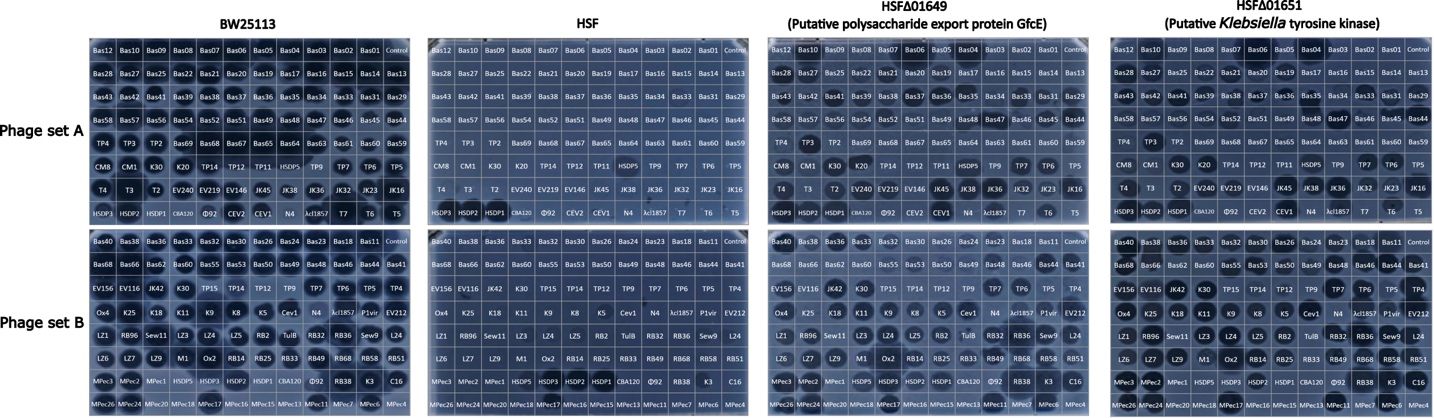

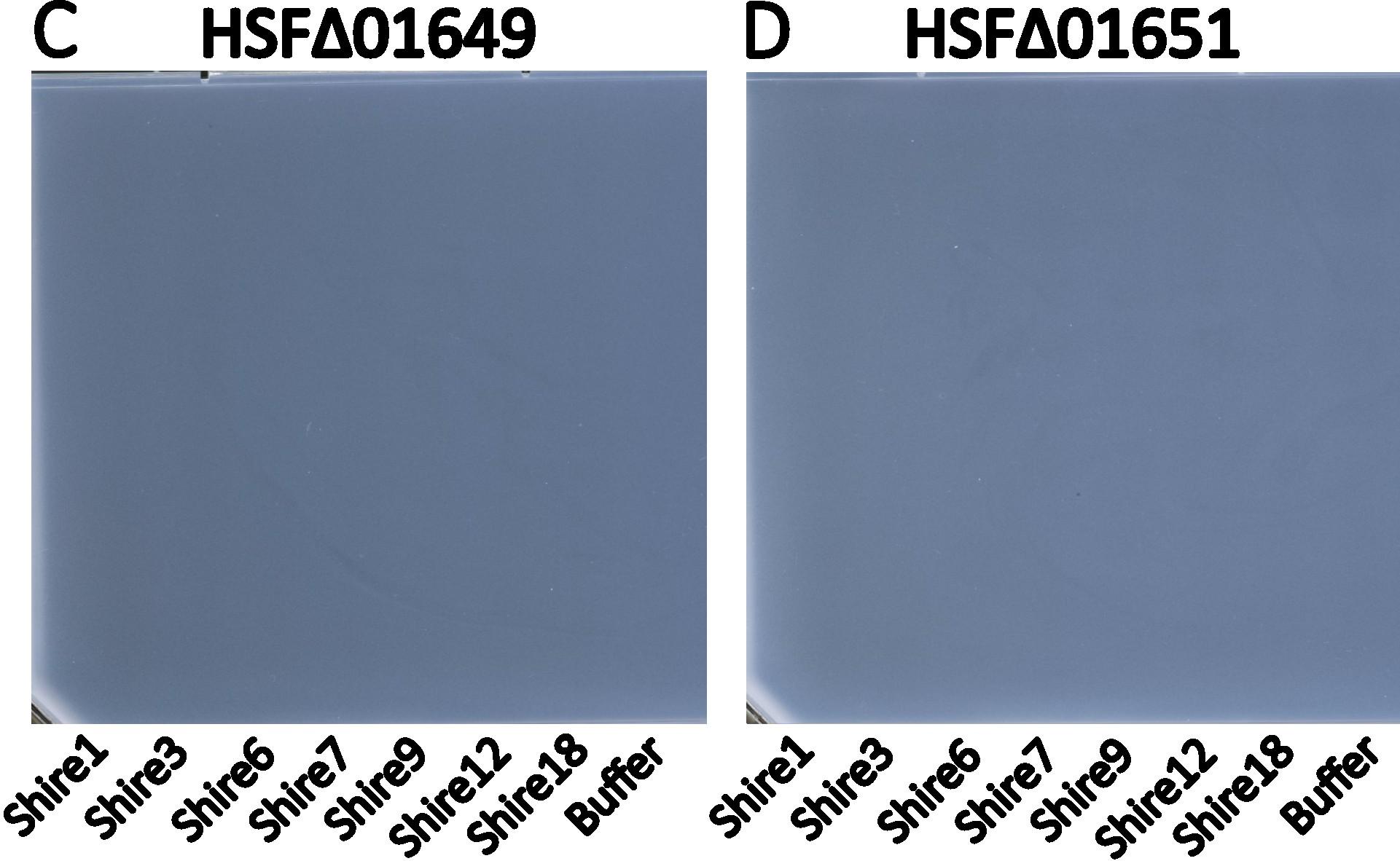

**Supplementary Figure S10. Host capsule expression in *E. coli* HSF restricts infection across a broad collection of 156 dsDNA phages.** Host susceptibility profiling demonstrating the protective barrier role of the HSF capsule. Diverse phage screening sets (A) and (B) composed of dsDNA phages capable of infecting the *E. coli* BW25113 reference strain largely fail to infect wild-type *E. coli* HSF strain. Conversely, the host becomes susceptible to a majority of these phages when capsule biosynthesis is disrupted via clean genetic deletions of the capsule-encoding genes, Δ01649 and Δ01651. Meanwhile, these same capsule-deletion strains remain entirely resistant to dsDNA environmental phages (C) and (D), which were originally isolated using *E. coli* HSF as the host. Titration of full panel of dsDNA phages on WT *E. coli* HSF is given in Supplementary Figure S4.

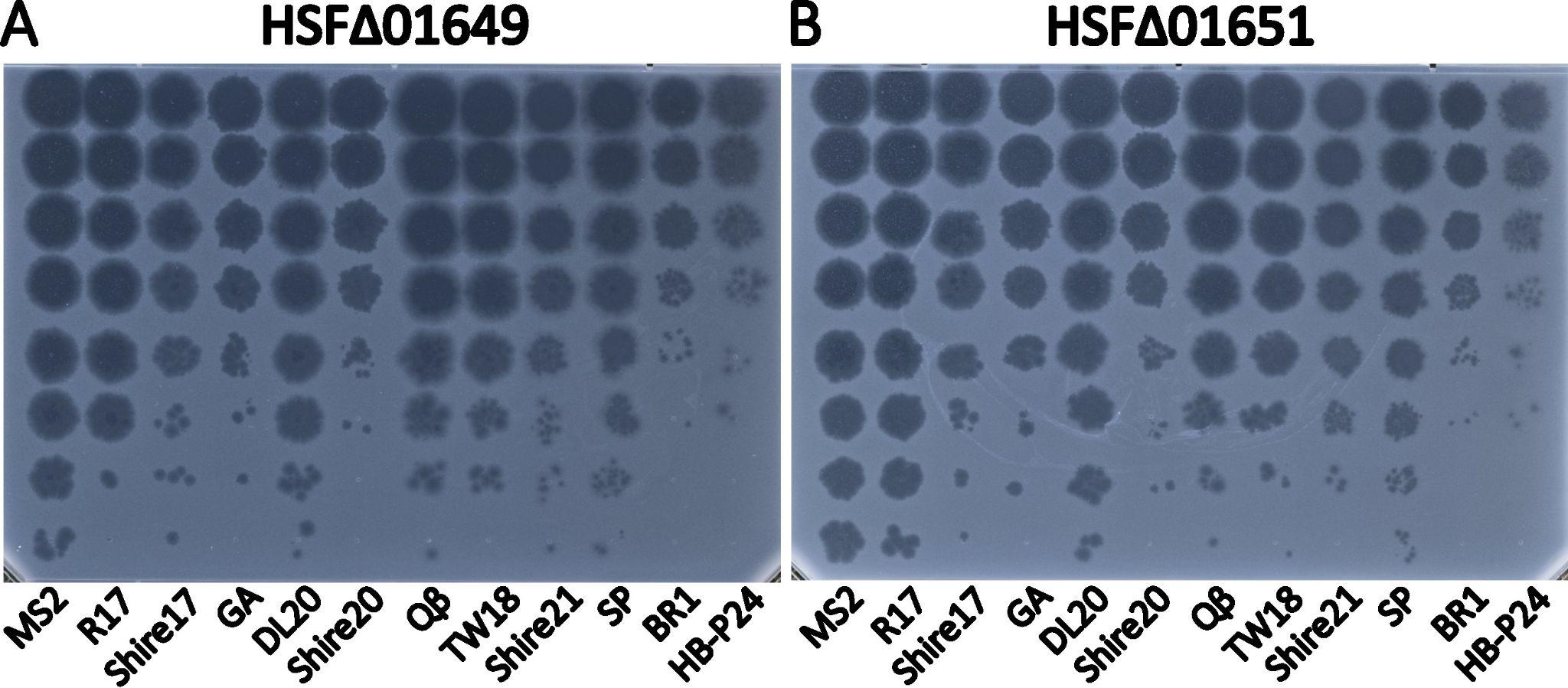

**Supplementary Figure S11. Capsule disruption does not impact ssRNA phage infection.** Host susceptibility profiling demonstrating that the capsular matrix does not obstruct ssRNA phage entry. Three representative ssRNA phage isolates selected from each distinct genogroup were titered via spot assays on (A) the *E. coli* HSF Δ01649 strain and (B) the HSF Δ01651 strain. Uninhibited plaque formation across both WT HSF and mutant backgrounds highlights that, unlike dsDNA phages, *Leviviricetes* infection pathways are unaffected by capsule.

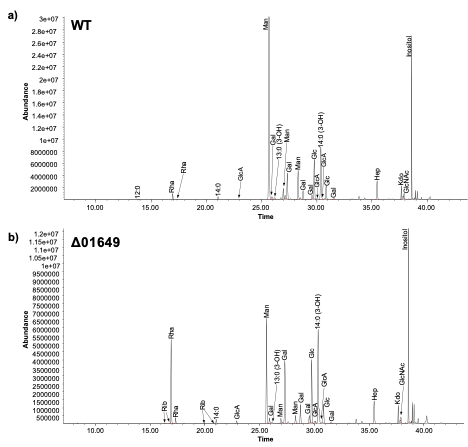

**Supplementary Figure S12. Chemical characterization of cell surface glycolipids isolated from wild-type *Escherichia coli* HSF (a) and the capsule-deficient *Δ01649* mutant (b)**. Deletion of the capsule gene (*01649*) decreased the relative abundance of mannose by half and increased the relative content of rhamnose and galactose in the *Δ01649* mutant. The loss of mannose was also accompanied by an increase in the relative abundance of fatty acids, reflecting a loss of a portion of the polysaccharide in the total analyzed pool of surface glycolipids. These results suggest that the parent strain is shielded by a mannose-rich polymer matrix of capsular polysaccharide. The details of compositional analysis can be found in Supplementary Tables S8 and S9.
